# An RNA Language Model trained on sequence alone reveals the structural logic of Internal Ribosome Entry Sites

**DOI:** 10.64898/2026.05.19.726202

**Authors:** Adam Sychla, Pierre Bongrand, Grant Yang, Aravind Iyer, Jacob Rulison, R. Alexander Wesselhoeft, Namita Bisaria, Silvi Rouskin

**Author notes:** These authors contributed equally (Adam Sychla, Pierre Bongrand). Senior author (Silvi Rouskin).

## Abstract

Millions of RNA sequences are readily available, but the structures that determine their function are not. Picornaviruses initiate translation through Internal Ribosome Entry Sites (IRESes), RNA elements that recruit ribosomes independent of the 5^′^ cap. These elements are large and highly divergent, and structural understanding remains limited to a handful of cases. We address this bottleneck by introducing an RNA language model (Albatross), trained purely on sequence, that predicts high-quality IRES structures at scale. We collect *in cellulo* chemical probing data for 96 full-length IRESes from divergent viruses and show that Albatross achieves far higher precision (0.80) than state-of-the-art predictions (0.47). Analyzing 75,000 IRES structures, we discover a novel Type II structural subclass and validate it experimentally. We demonstrate the pipeline broadly generalizes to identify functional structures, including tertiary contacts and alternative riboswitch structures. These findings establish Albatross as a scalable framework that accelerates RNA structure discovery and antiviral targeting.

## INTRODUCTION

RNA structures are known to mediate crucial cellular and viral functions, including translation, transcription, and immune response^1^. Predicting such structures from sequence would enable high-throughput characterization and study of complex host-viral interactions. Such functional RNA structures are highlighted by picornaviruses (*Picornaviridae*), a diverse and widespread family of RNA viruses with historic, ongoing, and emergent global health impacts, particularly among pediatric populations^2–8^. Members of this ubiquitous viral family are capable of infecting a wide variety of tissues, including neural, hepatic, cardiac, gastrointestinal, and respiratory, with outcomes ranging from asymptomatic infections to permanently debilitating diseases^9–13^. In addition to human disease, picornaviruses infect nearly all known vertebrates, posing a substantial risk of zoonotic spillover^14^. These viruses currently have high pandemic potential^15^ but no approved antiviral therapeutics^16^.

Picornaviruses typically encode a single polyprotein from their small (∼8 kb), positive-sense RNA genomes, with cap-independent translation initiation driven by large (*>*300 nt) structured RNA elements called Internal Ribosome Entry Sites (IRESes)^9^. IRESes are known to exhibit tissue and host-specific activities, helping determine spillover risk and severity of disease^10,17^. Beyond their biological significance, IRESes have found broad utility in biotechnology, with the EMCV IRES being a standard tool in bioengineering and gene therapy^18–20^. A more global understanding of rules governing IRES structure would provide paths toward improved pandemic readiness and inform the future design of targeted antivirals.

Although individual IRESes have been extensively studied, family-wide rules governing IRES function and tropism remain poorly understood. Massively parallel DNA synthesis enables high-throughput studies of short (∼200 nt) IRES fragments, but it cannot yet produce at scale the full-length sequences (400–1500 nt) needed to capture complete IRES structures. As a result, our understanding of full-length IRES architecture and function is largely limited to individual prototypical examples, and most IRES structures remain experimentally unresolved.

Thermodynamics-based RNA folding algorithms perform well on short sequences (*<*100nt) but require experimentally-derived constraints to obtain accurate structures for longer sequences such as IRESes^21–23^. Such models also become computationally intractable when accounting for many unknown nested, higher-order, or long-range interactions^24,25^. Meanwhile, predictive methods using machine learning have suffered from limited availability of high-quality RNA structure training data. Many models perform well on shorter sequences or in specific contexts but quickly fail when asked to extrapolate into new RNA families^23,26–28^.

Covariation analysis remains one of the best approaches for determining high-confidence RNA secondary structures^29–31^. However, these approaches depend on multiple sequence alignments of several closely related sequences with mutated and compensated base pairs to inform the analysis, which are often lacking, especially for highly divergent RNA elements like IRESes.

An additional difficulty in RNA structure analysis arises from the fact that any RNA will thermodynamically fold into some structure. However, it is unclear whether any of these substructures are functional or are simply a consequence of RNA’s tendency to base-pair. Systematic approaches have addressed this by requiring candidate structures to associate with a measured regulatory outcome^32^, which limits discovery to phenotypes that can be assayed at scale.

For this work, we sought to find broad, family-wide patterns in IRES activity and structure. Because precise IRES boundaries are often ill-defined, especially for divergent or understudied viruses, we generally use the term “IRES” throughout to refer to full picornaviral 5^′^ UTRs, while acknowledging that not all sequences within these regions necessarily contribute to translation initiation. For example, enteroviruses have a well-characterized cloverleaf structure at the 5^′^ terminus that is critical for genome replication rather than translation initiation^9,33^.

To address the structural bottleneck, we developed Albatross, an RNA Language Model obtained by continued pretraining of RiNALMo on ∼50,000 IRES sequences using only genomic RNA sequence data and a masked-token prediction objective^34^. Despite receiving no external knowledge of secondary structure, base pairing rules, or thermodynamics, the model learned an encoding of IRES structure from sequence alone^35^. We validated Albatross’s performance on a library of 96 full-length IRESes. Our *in cellulo* chemical probing data found that IRES structures were consistent across multiple cell types. Albatross independently recapitulated these experimental structures with very high precision, outperforming co-variation analysis (Infernal^30,31^), machine-learning based structure predictions (eFold^23^) and purely thermodynamic based models (RNAstructure)^21^.

In parallel, we measured translation driven by all 96 IRESes across six cell types, generating a comprehensive functional atlas of IRES activity. This revealed that the recently described Type V IRESes drive roughly twice the translational activity of the widely used EMCV IRES, and that many IRESes show significant cell-type-specific activity.

We probed Albatross’s mode of inference and found that it selectively predicts conserved, and thus likely functional structures rather than learning thermodynamics. This behavior allows us to prioritize predicted structures for experimental follow-up. We also generated high-quality structure maps for ∼75,000 IRESes available at http://www.albatrossrna.org. The large IRES structural data set allowed us to discover and validate a previously unreported divergent structural subclass of Type II IRES.

Finally, we show that this continued pretraining strategy improves with model size and generalizes beyond IRESes. Applying continued pretraining to hantavirus genome, we recovered known pseudo-circularization interactions; applying it to bacterial riboswtiches we identified conflicting base-pairing dependencies that reflected their two mutually exclusive functional states, that correspond to canonical forms. The model uncovered features such as tertiary contacts, pseudoknots, and alternative conformations, which are notoriously difficult for existing computational tools to predict. We thus propose Albatross approach as a general strategy for structural discovery across RNA biology.

## RESULTS

### Albatross predicts IRES structure from sequence alone

Studying IRESes as a comprehensive class remains a challenge, in large part limited by difficulties in accurate high throughput RNA structure measurement or prediction. To address this challenge, we instead sought to extract structure information directly from sequence. This is motivated by the fact that sequenced molecules must have survived evolutionary pressures and so implicitly as an aggregated whole they encode structural and functional constraints.

Recently, da Silva et. al demonstrated that dependency mapping is able to predict secondary structure of functional RNAs (*e.g.* tRNAs, RNAseP) using an Large Language Model (LLM) trained on sequences alone^35^. LLMs capture relations between tokens (base identity in the case of RNA) in their higher-dimensional encoding. Dependency mapping uses a trained LLM to measure how the prediction of a given nucleotide depends on a single point mutation elsewhere in the sequence, revealing a variety of structural elements. This approach is, in part, motivated by covariation analysis. If a mutation in one base leads to a changed prediction in another base, this indicates base pairing interactions (Figure 1A).

**Figure 1:**
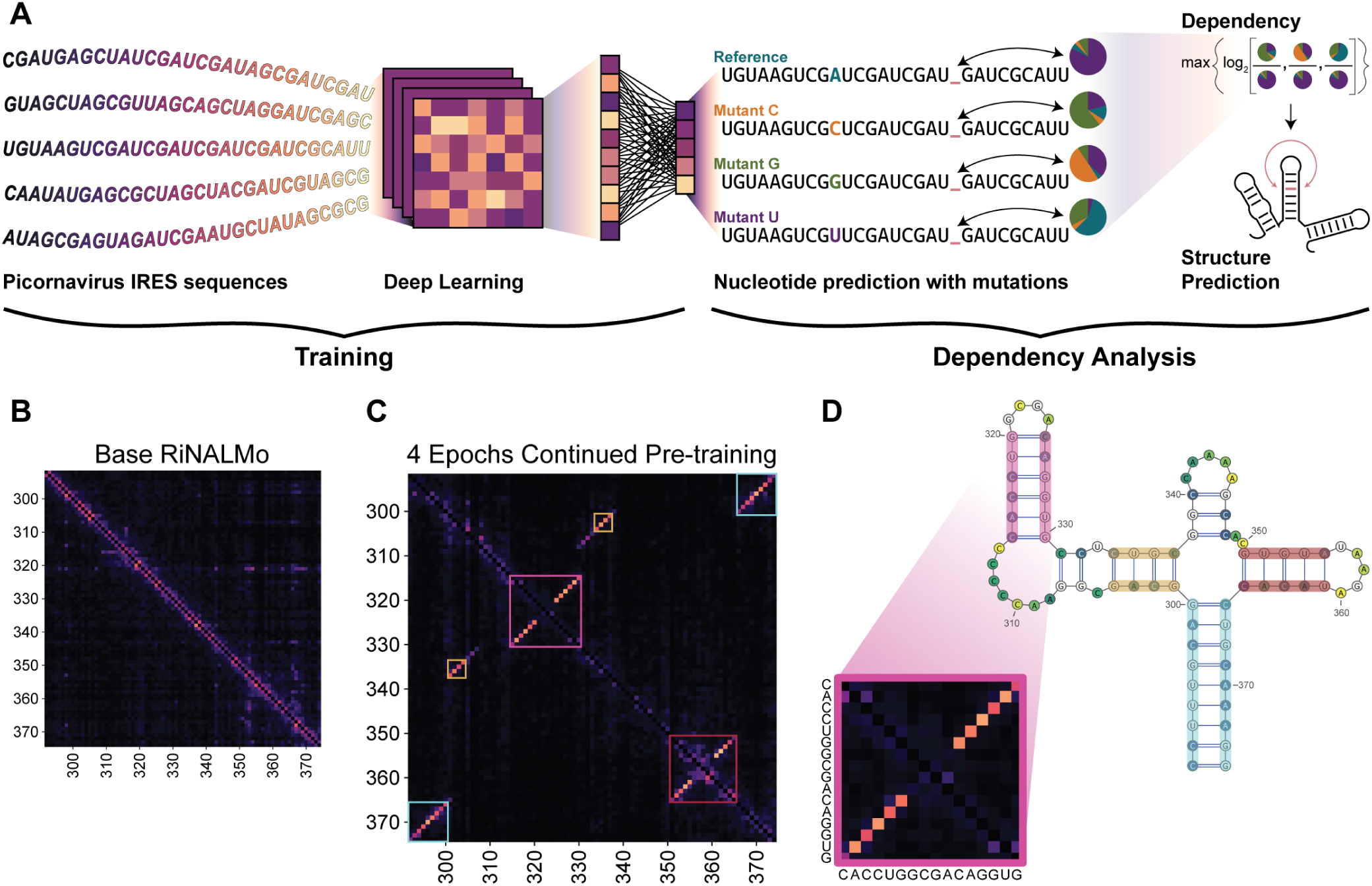
Albatross predicts IRES secondary structure from sequence alone. (**A**) Schematic of the dependency mapping approach. IRES sequences are used for continued pretraining of an RNA language model. For each position, the effect of every possible point mutation on nucleotide predictions is measured, generating a dependency map from which structure is inferred. (**B**) Dependency map of the EMCV IRES using the pre-trained model (before additional continued pretraining), showing no discernible structural signal. (**C**) Dependency map of the EMCV IRES after four epochs of continued pretraining. Antidiagonal features (dashed boxes) correspond to base-paired stems. (**D**) Albatross-predicted structure of the EMCV IRES overlaid on the experimentally validated secondary structure. Colors of the stems correspond to colored squares in C. Inset shows the dependency map for a stem region with corresponding sequence.

Several pre-trained RNA Language Models exist^34,36,37^. However, these models have been trained on extremely broad and divergent sequences and accordingly, performed poorly in our use case. For example, dependency mapping of the EMCV IRES returned noise (Figure 1B). Nonetheless, they capture some general patterns of RNA sequences internally so we decided to leverage this basis and continue pretraining an RNA Language Model on IRES sequences.

We started from RiNALMo, a 650M-parameter base model pre-trained on 36M non-coding RNA sequences^34^. Rather than training from scratch, we continued pretraining this model on IRES sequences, a transfer learning strategy widely employed in natural language processing and computer vision to adapt pre-trained models to new domains. We collected ∼10,000 picornaviral 5^′^ UTRs from the NCBI Virus Database^38^. For each, we performed a BLAST alignment against the nt database and collected the top 500 results^39^. After deduplication, this yielded ∼200,000 IRESes for continued pretraining of RiNALMo. Importantly, training was done exclusively on RNA sequences with no explicit information on structure or base pairing rules. We dubbed the resulting model Albatross.

Dependency mapping of the EMCV IRES using Albatross revealed antidiagonal dependencies corresponding to stems in the experimentally validated secondary structure (Figure 1C, D)^40,41^.

### Experimentally derived IRES structures are stable across cell types

While Albatross was able to predict a well-studied canonical structure, we wanted to test it against an expanded range of ground truth structures. We generated a library of 96 IRESes from a range of hosts and IRES structural types (Figure 2A)^42^. Approximately half of the IRESes come from viruses with human as the primary host, while the rest span vertebrate hosts including amphibians, ungulates, and non-human primates. Each construct consisted of a barcode, IRES, and firefly luciferase. We procured a final mixed RNA library in both linear and circRNA formats.

**Figure 2:**
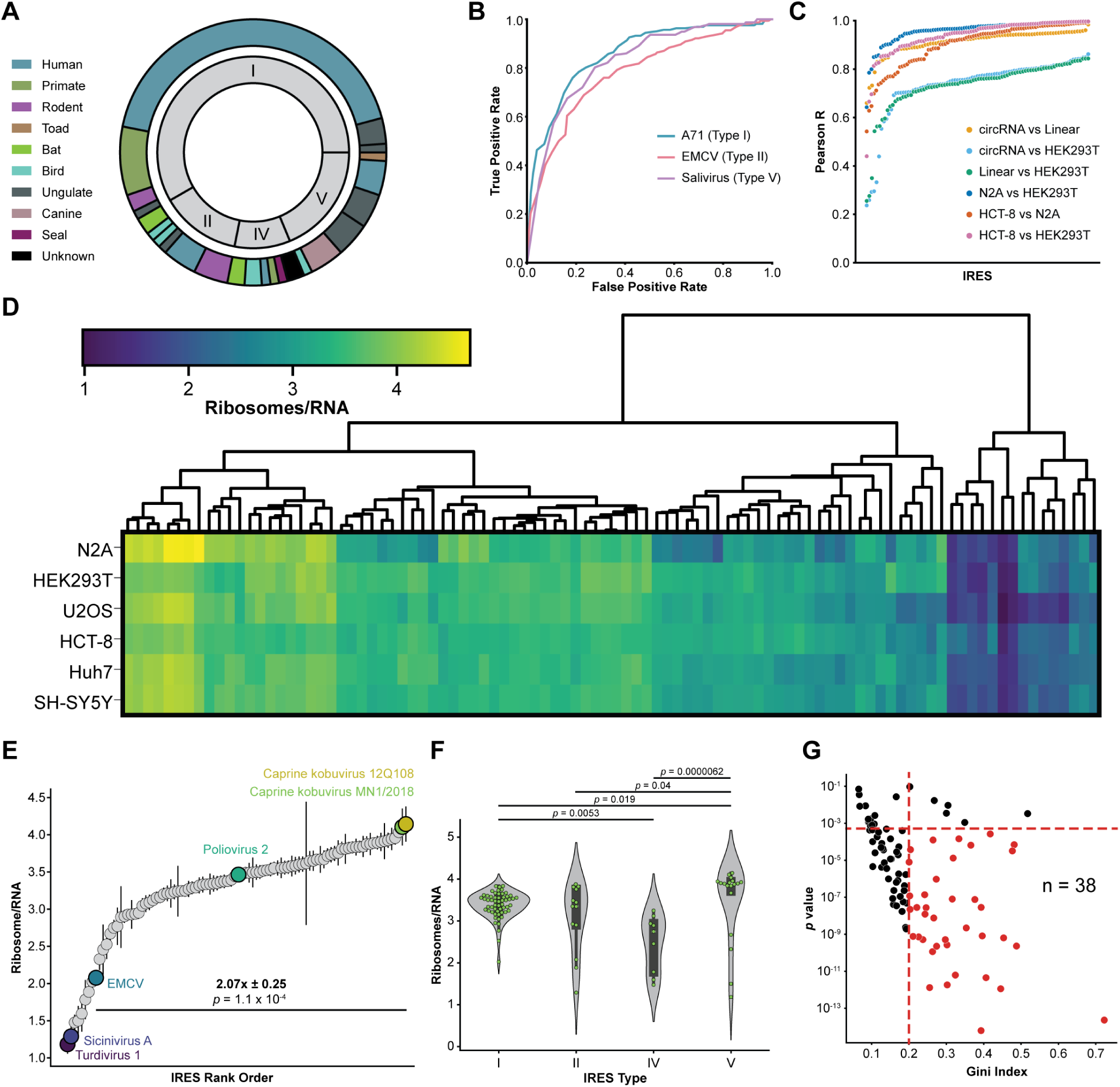
Experimental structure-function altas of 96 IRESes. (**A**) Library composition by host (outer) and IRES type (inner). (**B**) DMS-MaPseq predicted structure in HEK293T cells AUC-ROC for three ground truth literature IRES structures. (**C**) Pearson R/Construct between DMS-MaPseq experimental replicates and conditions. circRNA and linearRNA refer to the in vitro probed library; labels naming a cell type refer to the same library probed in cells. (**D**) Heatmap of ribosomes/RNA for all 96 IRESes across six cell types, with hierarchical clustering. Color scale indicates ribosomes/RNA. (n = 4/cell type). (**E**) IRES activity (ribosomes/RNA) ranked across 96 IRESes in HEK293T cells. Select IRESes are labeled. EMCV activity relative to the top-performing Caprine kobuvirus 12Q108 is indicated. *p*-values from Student’s t-test, n = 4 replicates. (**F**) Ribosomes/RNA grouped by IRES structural type. *p*-values from Kruskal-Wallis H test with Dunn post-hoc analysis and Bonferroni correction, n = 4 replicates. (**G**) Tropism significance (p-value) versus effect size (Gini Index) for each IRES. Dashed line indicates significance threshold after Bonferroni correction. 38 IRESes with moderate to strong tropism. *p*-values from ANOVA.

We used DMS-MaPseq to generate experimentally constrained *in vitro* and *in cellulo* structure models for our library^43–45^. Our Enterovirus A71, EMCV, and Salivirus models showed high agreement with published structures (Figure 2B)^40^. We found that IRES structure generally exhibited minimal conformational differences between circular and linear topologies and across three cell types, with the biggest differences found between *in cellulo* and *in vitro* refolded RNAs (Figure 2C). Given the high concordance between cell types, we proceeded with the HEK293T *in cellulo* structures as our “ground truth”, where we had sufficient coverage on 95 sequences. We provide DMS-MaPseq constrained structure models for all 95 IRESes at www.rnandria.org^23^.

The same set of barcoded constructs used to generate this structural ground truth also let us measure IRES translational output, providing a matched dataset for asking how structure relates to function.

### Type V IRES activity doubles that of standard biotech IRESes

Our IRES library was among the largest reported for full-length IRESes, spanning 365 nt to 1431 nt, in contrast to previous studies that screened ∼200 nt or smaller IRES fragments^46–48^. This provided us an opportunity to create the largest public IRES activity atlas.

IRES activity is traditionally screened for using dual reporter assays. However, these assays are confounded by multiple sources of false positives. The test sequence may harbor cryptic promoters or splice sites that generate monocistronic transcripts of the downstream reporter and translational readthrough ranges from 0.1 to 1.5%^49–52^. The circular architecture of our library overcomes these challenges by containing a single ORF driven by the IRES and no 5^′^ end.

We used polysome profiling to characterize the activity of our entire library in a single experiment^53^. Our optimized protocol enabled us to consistently recover highly resolved polysome fractions up to heptasomes across a variety of cell types, with high reproducibility across biological and technical replicates (Figure S3).

We profiled IRES activity in a total of six cell lines representing a range of tissues, namely HEK293T (renal), HCT-8 (colorectal), Huh7 (hepatic), N2A (neural murine), SH-SY5Y (neural), and U2OS (osteal) (Figure 2D, S1, S2, S3). The cell lines in these experiments represent a wide range of tissues, most of which are relevant in picornaviral infections. Crucially, this atlas is comprised of identical experimental, collection, and analysis procedures across the cell types ensuring the data is directly comparable and representative of cell type specific effects.

In HEK293Ts IRES activity spanned approximately half an order of magnitude (Figure 2E). Notably, the EMCV IRES, typically used in synthetic constructs and gene therapy designs, exhibited comparatively poor performance (87^th^/96). Surprisingly, the top ten expressing IRESes in HEK293Ts were all Type V, from viruses with either ungulate or canine natural hosts. Type V is the most recently classified IRES, first identified in 2011, and remains the least studied of the five types, with only a handful of individual members functionally characterized to date^54^.

To confirm that the polysome profiling data corresponded to an alternative method of translational readout we turned to a classic dual luciferase reporter assay (Figure S4). We tested the top two and bottom two performing IRESes, the median performing IRES (Poliovirus 2), and the EMCV IRES. All IRESes corresponded to the rank order from polysome profiling (Figure S4). Beyond rank order the two assays recapitulated relative expression (*e.g.* Caprine kobovirus 12Q108 had 2.07 ± 0.25 and 2.26 ± 0.25 fold activity compared to EMCV in polysome profiling and dual luciferase respectively).

Type V’s advantage was not limited to HEK293Ts: across the full panel of cell lines, Type V IRESes matched or exceeded the other structural types in our library in every condition tested (Figure 2F, S5), raising the possibility that Type V’s unique architecture leads to higher translational output. On the other hand, Type IV IRESes showed the opposite pattern, consistently ranking lowest across all six cell types (Figure S5).

### IRESes exhibit moderate tissue-specific activity

On a broad scale, the rank order of IRES activity was preserved between cell lines (Figure 2D), consistent with the common translation factors shared across these cell types and the classification of IRESes into a limited number of large-scale structural types that serve as central scaffolds across many hosts and tissues^9^. Nonetheless, IRESes that deviated from such ordering were identifiable in every cell comparison.

We found that 77/96 IRESes displayed significant tropism (*i.e.* cell type specific activity) (Figure 2G). Using Gini Index (a measure of statistical dispersion) as a measure of effect size, we found 38 of these (∼40% of the total) exhibited moderate tropism(Figure 2G). For example, Rhimavirus A had a 1.9 ± 0.1 fold expression (N2A/U2OS) and Seal Picornavirus type 1 had 1.6 ± 0.2 fold expression (293T/U2OS). Whereas, Caprine kobovirus 12Q108 (Type V) had barely any tropism at 1.2 ± 0.2 fold expression (293T/HCT-8).

At the extreme ends of expression within human cells, which matter most for RNA therapeutics^19,47^ and viral host barriers^55^, IRES type was the strongest predictor of activity (Figure S5). Yet new picornaviruses are sequenced and deposited much faster than their IRESes can be structurally characterized^38^ We wanted to capture the full spectrum of structural diversity within and across types among these sequences. Chemical probing is laborious, slow, and impractical for thousands of IRESes. The ideal is an equally good approach that is fast and cheap and can accurately predict the structure, and therefore inform the function, directly from sequence.

### Albatross predicts diverse IRES structures and complex tertiary contacts

To test the performance of Albatross on structure prediction from sequence, we used our 95 HEK293T DMS-MaPseq constrained structure models as “ground truth”. While these models are imperfect representations of reality, no other large-scale dataset of full-length IRES structures exists and the current state-of-the-art for mapping full-length IRESes remains chemical probing^40,48,56^. Importantly, all of our models had AUC-ROC values between 0.79-0.93, indicating high agreement between the predicted structures and the underlying DMS reactivity data (Figure S6)^57,58^.

From the dependency maps, we generated explicit structures by applying a binary filter to assign base pairing (Methods). Our filtering did not specify base pairing rules, allowing for non-canonical base pairs. Crucially, Albatross never had access to structural data before or while it made a prediction. All analysis followed from the output dependency maps. Because no structural information entered training, whether a benchmark sequence appeared in the training set has no bearing on the model’s access to its structure. Sequence overlap between training and evaluation sets therefore does not confound this comparison.

We postulated that a greater training set could improve performance so we developed a larger training set, prioritizing the expansion of small clusters for a total of ∼500,000 sequences. We compared the microF1 over 10 epochs of training (Figure 3A). While the 500k training set performed better than 200k, both rapidly reached a peak microF1 and worsened with longer training. The surprising degradation of performance indicated some form of over-training was occurring. We speculated that with more learning steps Albatross had worse scores through two mechanisms: 1) Albatross over-optimized for the highly represented sequences leading to worse performance on lowly represented sequences; 2) Albatross starts to forget its pretraining as we overtrain on these sequences^59^.

**Figure 3:**
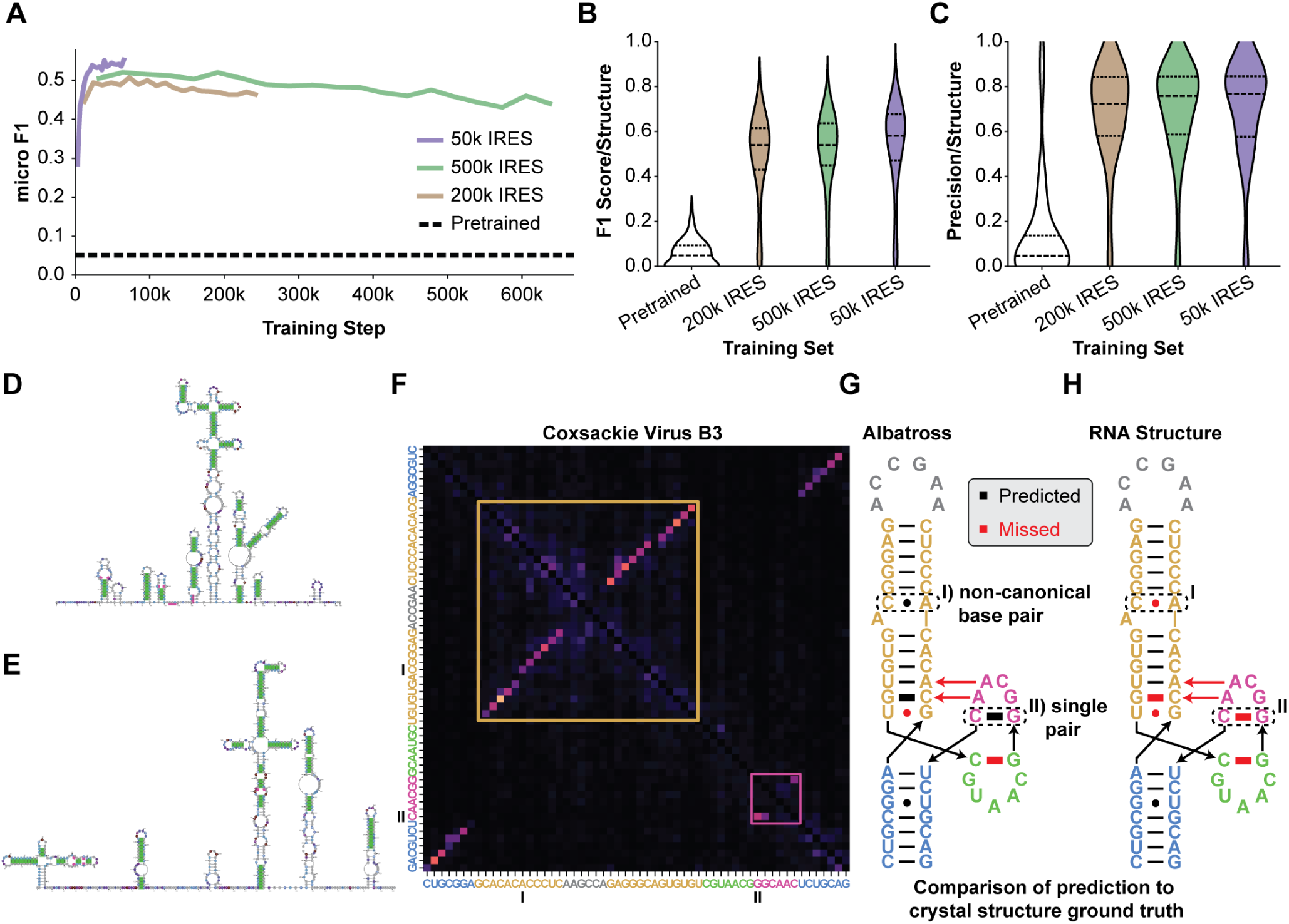
Albatross predicts IRES structure with high precision. (**A**) MicroF1 over training steps for the 200k, 500k, and 50k training sets. Dashed line, pre-trained baseline. (**B**) F1 score per structure across training sets. (**C**) Precision per structure across training sets. (**D, E**) Representative IRES secondary structures for EMCV (Type II) and Enterovirus A71 (Type I). Base pairs predicted by Albatross that agree with the consensus literature structures are colored as true positives (green), while Albatross predictions absent from the consensus literature structures are colored as false positives (pink). (**F**) Albatross dependency map of the Coxsackievirus B3 eIF4G-binding Domain V. (**G**) Albatross-predicted base pairs compared to the crystal structure, with predicted interactions in black and missed interactions in red. Albatross correctly identifies (I) a non-canonical A • C pair and (II) an isolated single G • C stabilizing interaction. (**H**) RNAstructure Fold thermodynamic prediction for the same region.

We considered if a smaller but more diverse training set could recapitulate the “ground truth” structures better. We truncated our 500k training set down to ∼50,000 sequences, prioritizing balanced inclusion from each *>*70% similarity cluster (Methods)^60^. This smaller library attained a substantially better microF1 than the two larger libraries (Figure 3B). In addition to improved performance, the smaller data set required less training, taking only twelve hours on one GPU. We evaluated F1 for each individual IRES using the best microF1 checkpoint for each training set. F1 scores trended upwards from the 200k to 500k to 50k training sets (Figure 3A). The best structure, for WUHARV Enterovirus 2, a Type I IRES, had an F1 of 0.89.

Across the 95 IRESes, Albatross achieved consistently high precision, with a median of 0.80 and individual structures reaching as high as 1.0 (Figure 3C, S7, Table 1)^40,41,61,62^. This means that when Albatross predicts a base pair, that pair is usually present in the experimental data. We decided to move forward with the 50k Albatross model for the remainder of the paper.

**Table 1.** Model Performances.

| Model | F1<br>Mean | F1<br>Median | Precision<br>Mean | Precision<br>Median | Recall<br>Mean | Recall<br>Median |
| --- | --- | --- | --- | --- | --- | --- |
| pretrained | 0.06 ± 0.06 | 0.05 | 0.14 ± 0.24 | 0.05 | 0.06 ± 0.06 | 0.05 |
| Albatross 200k | 0.48 ± 0.20 | 0.54 | 0.67 ± 0.23 | 0.72 | 0.39 ± 0.18 | 0.44 |
| Albatross 500k | 0.50 ± 0.18 | 0.54 | 0.68 ± 0.23 | 0.76 | 0.41 ± 0.16 | 0.43 |
| Albatross 50k | **0.54 ± 0.21** | **0.60** | **0.73 ± 0.22** | **0.80** | **0.44 ± 0.19** | **0.48** |

While family wide structures for IRESes are unavailable, specific examples have been well studied. We compared the Albatross predictions to the literature structures of EMCV (Type II), A71 (Type I), and Salivirus (Type V) (Figure 3D, E, S7)^40,41,62,63^. In each case Albatross successfully predicted most stems and crucially did so with few false positive base pairs.

There are no full-length crystal structures of picornaviral IRESes but several fragments have been resolved. We compared the Albatross-derived dependency map of the Coxsackievirus B3 IRES against the recently solved crystal structure of the eIF4G-binding domain V^64^. Albatross correctly predicted 19 of 21 base pair interactions, outperforming the thermodynamics-based RNAstructure algorithm, which successfully identified only 16 interactions (Figure 3F–H). Notably, Albatross independently predicted both a non-canonical A • C pair and an isolated single G • C base pair stabilizing interaction first revealed by the crystal structure (Figure 3G, H). These findings demonstrate the potential for dependency maps to identify base interactions missed by conventional RNA secondary structure prediction approaches.

### Albatross achieves dramatically higher base-pair precision than state-of-the-art RNA structure prediction methods

We wanted to compare Albatross with state-of-the-art analysis methods. To that end, we chose Infernal with CaCoFold to represent covariation analysis, eFold to represent machine intelligence methods, and RNAstructure to represent thermodynamic prediction^21,23,30,31^. We also tested using the Albatross predicted base pairs as hard constraints on RNAstructure’s predictions. The “ground truth” data uses DMS reactivity to constrain RNAstructure giving the RNAstructure method an inherent advantage in this comparison. For Infernal, we used Rfam’s curated family alignments and therefore represents the strongest available covariation-derived structure for these families.

We compared F1, precision, and recall for each IRES (Figure 4A–C, Table 2). On F1, Albatross outperforms both Infernal and eFold and was comparable to RNAstructure alone, while Albatross-constrained RNAstructure exceeds every other method (median F1 0.64; Figure 4A). Albatross alone achieved much higher precision than all other methods, with a median of 0.80 against 0.47 for Infernal, 0.44 for eFold, and 0.47 for RNAstructure, and it occupied its own statistical group (Figure 4B). Its recall was not significantly different from that of Infernal or eFold (Figure 4C). All Albatross values use the cross-validated filter threshold (Methods).

**Figure 4:**
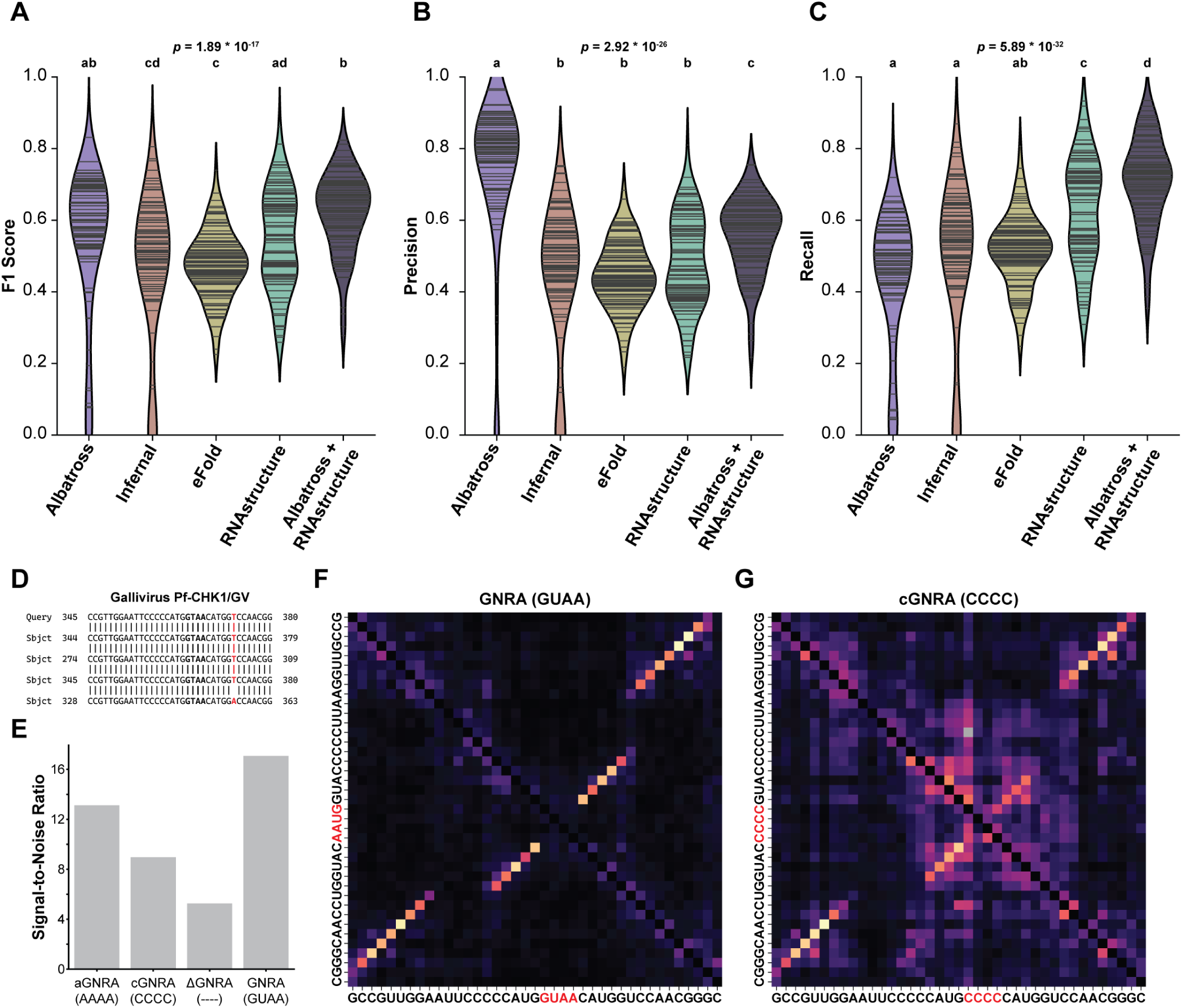
Albatross outperforms state-of-the-art prediction methods on IRESes. (**A**) Violin plots comparing per-structure F1 scores between Albatross, Infernal, eFold, RNAstructure, and Albatross constraining RNAstructure predictions (*p* = 1.20 × 10^−14^ by Welch’s ANOVA, grouped by Dunn Post-Hoc Test with Holm method adjustment, n = 95 IRESes). (**B**) Violin plots comparing per-structure precision between Albatross, Infernal, eFold, RNAstructure, and Albatross constraining RNAstructure predictions (*p* = 1.21 × 10^−26^ by Welch’s ANOVA, grouped by Dunn Post-Hoc Test with Holm method adjustment, n = 95 IRESes). (**C**) Violin plots comparing per-structure recall between Albatross, Infernal, eFold, RNAstructure, and Albatross constraining RNAstructure predictions (*p* = 5.74 × 10^−24^ by Welch’s ANOVA, grouped by Dunn Post-Hoc Test with Holm method adjustment, n = 95 IRESes). (**D**) Per-IRES comparison of precision, colored as in (**B**): Albatross scored higher for 87 IRESes, Infernal for 4. (**D**) BLAST alignment of the Gallivirus Pf-CHK1/GV GNRA tetraloop region (bold) against the Albatross training set. Only four related sequences exist, with the divergent nucleotide in each marked in red. (**E**) Signal-to-noise ratio of the GNRA stem signal in dependency maps of the Gallivirus IRES bearing different tetraloop sequences: aGNRA (AAAA), cGNRA (CCCC), ΔGNRA (loop deleted), and wild-type GNRA (GUAA). (**F,G**) Dependency maps of the Gallivirus IRES with the wild-type GNRA tetraloop (GUAA, **G**) or the CCCC substitution (**H**).

**Table 2.** Comparison of RNA structure prediction models.

|  | Albatross<br>(50k) | Infernal | eFold | RNAstructure | Albatross +<br>RNAstructure |
| --- | --- | --- | --- | --- | --- |
| F1 Mean | 0.54 ± 0.21 | 0.46 ± 0.21 | 0.48 ± 0.10 | 0.55 ± 0.14 | **0.62 ± 0.11** |
| F1 Median | 0.60 | 0.51 | 0.48 | 0.55 | **0.64** |
| Precision Mean | **0.73 ± 0.22** | 0.44 ± 0.20 | 0.45 ± 0.09 | 0.48 ± 0.13 | 0.55 ± 0.11 |
| Precision Median | **0.80** | 0.47 | 0.44 | 0.47 | 0.58 |
| Recall Mean | 0.44 ± 0.19 | 0.49 ± 0.22 | 0.51 ± 0.10 | 0.63 ± 0.15 | **0.71 ± 0.12** |
| Recall Median | 0.48 | 0.54 | 0.51 | 0.66 | **0.73** |

These data position Albatross as complementary to thermodynamic prediction rather than as a replacement for it. Thermodynamic models score every possible structure by free energy minimization and return the most stable one, so they recover a larger fraction of the true pairs but also call many pairs the RNA does not form. Albatross calls a smaller set but far more reliably, because it reports only the pairings that are constrained within a given family sequence variation. Combining the two methods therefore captures both strengths: supplying Albatross pairs as hard constraints raises RNAstructure’s median precision from 0.47 to 0.58 and its median F1 from 0.55 to 0.64, with recall rising from 0.66 to 0.73, the highest of any method tested (Figure 4A–C).

### Albatross uses sequence context to complement covariation data in its predictions

Given that dependency mapping is conceptually similar to covariation analysis we wanted to know how Albatross was able to correctly predict base pairs that covariation was not. One of the most striking cases was a highly divergent Gallivirus IRES whose GNRA elbow structure aligned to only four sequences in the training data, with a single mutation among them (Figure 4D)^65^. With so few related sequences, covariation analysis is effectively impossible, and accordingly Infernal failed to predict this structure. Yet Albatross predicted it correctly.

If Albatross cannot rely on covariation for the Gallivirus IRES, it must be using a different strategy. We hypothesized that the model had learned to associate specific sequence motifs with structural outcomes, namely that GNRA tetraloops are consistently followed by elbow structures. To test this, we performed dependency mapping on the Gallivirus IRES with mutations in the GNRA motif: substitution to all As, all Cs, or complete deletion (Figure 4E–G, S8). The wild-type sequence produced clear dependencies corresponding to the elbow. Even a conservative GU to AA substitution introduced substantial noise in the GNRA stem, and substitution to CCCC further increased noise that propagated into the second stem of the elbow (Figure 4E, H, S8). Deletion of the GNRA motif eliminated the stem entirely, confirming that both motif identity and spacing drive the prediction (Figure 4E, S8). These results demonstrate that Albatross has learned to associate sequence motifs with structural context, a capability that extends beyond what covariation analysis can access.

### Thermodynamic stability is not sufficient for structure prediction

For the Gallivirus mutants, our filtering protocol could still reconstruct the correct structure despite this noise. This led us to investigate the underlying logic used by Albatross to drive its predictions. If Albatross relies on fundamental physical principles (*e.g.* thermodynamics), it should predict most structural features of the target RNA. If instead, the model predictions are driven by sequence context, its predictions would be focused on common structure features learned from examples in the training set. For that to occur the predicted structures have to be evolutionarily maintained in the training data, which would imply they provide a functional benefit to the virus. Fundamentally, the model’s mode of reasoning changes the interpretation of the predicted signals.

We trained a smaller, more focused model on ∼8,500 A71 IRES sequences. We identified two equal length regions in the A71 sequence that were predicted as either structured (12-6 hairpin loop) or unstructured by the model and in the literature (Figure 5A, S9). We designed a synthetic sequence that forms a 12 base pair hairpin (by thermodynamics) and did not appear anywhere in the training set (Figure S9). If the model’s predictions are driven by thermodynamics, replacing the unstructured region with the synthetic stem should predict the hairpin. However, no structure was predicted (Figure 5B). The naturally structured region was unaffected.

**Figure 5:**
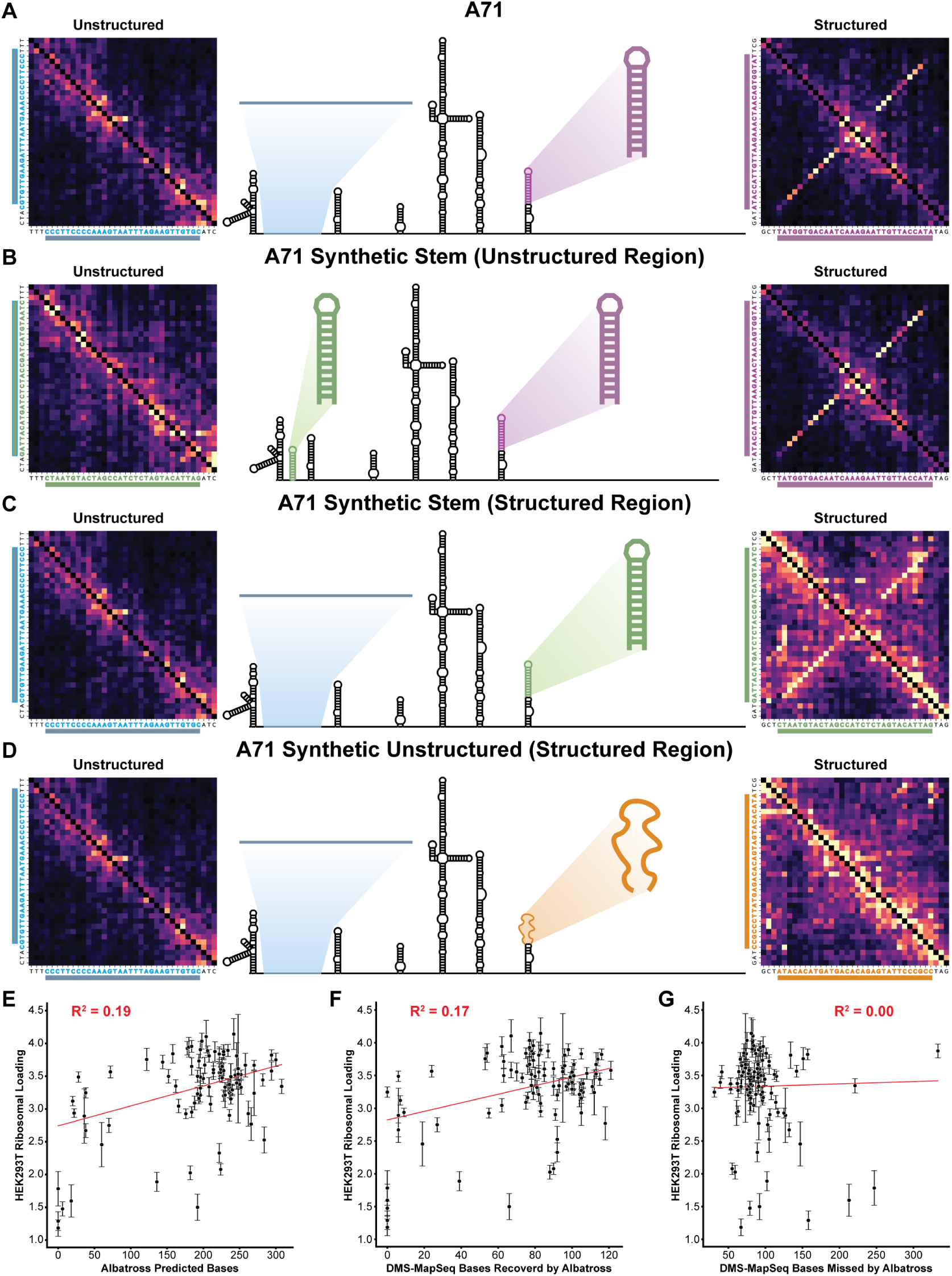
Albatross predicts a context-dependent subset of base pairs that is associated with IRES activity. (**A–D**) Dependency maps for the natively unstructured (left) and natively structured (right) region of the A71 IRES; central schematic highlights both regions and the secondary structure. Synthetic sequences shown in Figure S9. (**A**) Native A71. (**B**) A thermodynamically stable synthetic stem (green) in the unstructured region: no dependency map signal. (**C**) The same stem in the structured region: dependency signal observed. (**D**) A synthetic sequence of low thermodynamic stability (orange) in the structured region: no signal. (**E**) Albatross-predicted paired nucleotides per IRES versus ribosomal loading in HEK293T cells. (**F, G**) DMS-MaPseq pairs recovered by Albatross (**F**) and missed by Albatross (**G**), against ribosomal loading. Red line, linear fit; *R*^2^ and error bars, SD of n = 4.

These data show that the model does not simply apply generalized thermodynamic rules, corroborating the negative data from IRES predictions using the base RiNALMo (Figure 1B). Therefore dependency mapping cannot be broadly used on base models when investigating new RNA classes. Instead, continued pretraining, as in this work, is required.

We then took the same synthetic hairpin sequence and replaced the naturally structured region (Figure 5C). Even though the same sequence was being analyzed, in this context, the model correctly predicts the structure. This supports the premise that Albatross learns structural conservation that depends on sequence context.

Though Albatross does not apply generalized thermodynamic rules in its predictions, we wanted to investigate whether the model understood the physical constraints within its predictions. Earlier, we demonstrated that the model predicts non-canonical base pairs so it is not strictly held to Watson–Crick–Franklin base pairing rules (Figure 3G). However, the predicted A • C pairing in that example is physically possible^64^.

We designed another synthetic sequence of equal length to the hairpin but deliberately unstructured (Figure S9). When replacing the natural hairpin with this sequence, Albatross no longer predicted a stem indicating the model has internalized some physical constraints on pairing (Figure 5D).

These data altogether indicate that thermodynamic stability alone is not sufficient for prediction and dependency maps report spatial and evolutionary coupling rather than base pairing. Albatross identifies nucleotides whose identities constrain one another, which are typically proximal in three-dimensional space, including canonical helices (Figures 1D, 3D, 3E) and non-canonical pairs (Figure 3G).

### Albatross predicts a subset of base pairs associated with IRES activity

RNA’s propensity to base-pair generates many stable substructures that are not maintained by selection, and it is difficult to separate them from functional ones. To ask whether the pairs Albatross retains carry functional information, we compared predicted structure content against ribosomal loading in HEK293T cells. The number of Albatross-predicted paired nucleotides per IRES correlated with ribosomal loading (R^2^ = 0.19), as did the number of predicted stems (R^2^ = 0.20) (Figure 5E, S10A). The same metrics computed on the DMS-MaPseq constrained structures showed little relationship (R^2^ = 0.12, 0.10 respectively) (Figure S10B, C), similar to thermodynamic predictions from RNAstructure (R^2^ = 0.12, 0.11 respectively) (Figure S10D, E).

Because the dependency map predicted pairs are a subset of those supported by chemical probing, we partitioned each measured structure by whether Albatross recovered the pair. Pairs recovered by Albatross tracked ribosomal loading (R^2^ = 0.17), while pairs the model missed did not (R^2^ = 0.00) (Figure 5F, G). The correlation is modest, as expected for a coarse metric of structure tested against a single function measured in cultured cells. It is nonetheless specific to the pairs Albatross recovered.

*in cellulo* chemical probing reports all base pairing an RNA adopts, both elements maintained by selection and pairing that forms because the sequence permits it. In contrast, Albatross selectively predicts structures that are likely to carry function.

### Dependency mapping at scale identifies a novel Type II structural subclass

We then sought to analyze picornaviral IRESes as a class. We ran ∼75,000 unique IRES sequences through Albatross, generating a dependency map for each and manually labeling them by IRES type. These maps and predicted structures are freely available at www.albatrossrna.org.

We sought to analyze the structure predictions as a collective family of structures. To that end, utilized a modified alignment algorithm on the resulting dependency maps (Methods). In short, each dependency map was transformed, mapping each antidiagonal of the *n x n* map to a column of a *n x* (2*n*−1) matrix. An adjusted Needleman–Wunsch algorithm aligns corresponding stems and provides a similarity score (Figure 6A).

**Figure 6:**
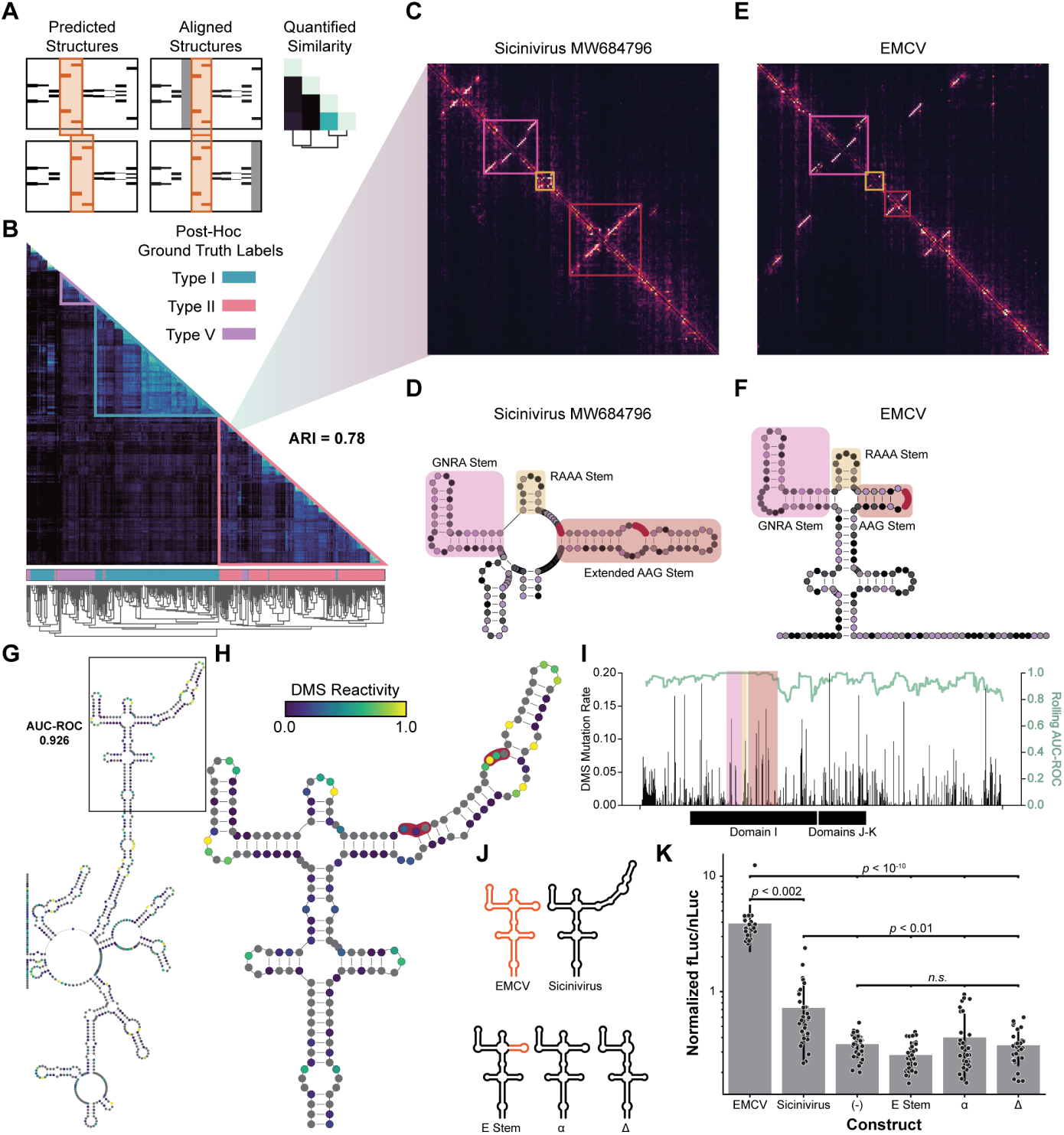
Structural clustering across a large IRES dataset reveals an extended stem in a Type II subclass. (**A**) Schematic of the dependency-map alignment procedure: predicted structures (left) are aligned to one another with gaps inserted as needed (middle), yielding pairwise similarity scores (right) used for hierarchical clustering. (**B**) Hierarchical clustering of alignment similarity scores across ∼75,000 predicted IRES structures. Colored bars below the matrix show post-hoc manual type labels not available to the unsupervised clustering(Type I, teal; Type II, pink; Type V, purple), demonstrating strong agreeement. (**C**) Dependency map of the Sicinivirus MW684796 IRES, with the GNRA stem (magenta), RAAA stem (gold), and extended AAG stem (red) boxed. (**D**) Secondary structure of the Sicinivirus MW684796 IRES Domain I, with the same three stems highlighted. Red circles mark the two conserved AAG motifs within the extended stem. (**E**) Dependency map of the canonical Type II EMCV IRES, boxed as in (**C**) for comparison. (**F**) Secondary structure of the EMCV IRES Domain I, highlighted as in (**D**). EMCV contains a single AAG motif rather than the extended, two-motif stem seen in Sicinivirus MW684796. (**G**) DMS-MaPseq validation of the Albatross-predicted Sicinivirus MW684796 IRES structure. Nucleotides are colored by DMS reactivity (scale in **H**); agreement between predicted base-pairing and reactivity across the full IRES is AUC-ROC = 0.926. Boxed region is enlarged in (**H**). (**H**) Enlarged view of the boxed Domain I region in (**G**). Red-outlined nucleotides mark the two conserved AAG motifs within the extended stem. (**I**) DMS mutation rate (bars, left axis) and rolling-window AUC-ROC (green line, right axis) across the Sicinivirus MW684796 IRES. Shaded regions mark the GNRA (pink), RAAA (gold), and extended AAG (red) stems as in (**C–F**); black bars below indicate the annotated Domain I and Domains J-K regions. (**J**) Visual summary of Sicinivirus MW684796 mutants screened in dual luciferase assay. (**K**) Dual luciferase assay of Sicinivirus MW684796 mutants in UMNSAH/DF1 cells. *p*-values from Welch’s ANOVA with Dunn Post-Hoc Test with Holm method adjustment.

Hierarchical clustering on these alignments naturally groups IRESes by type without pre-labeling (Figure 6B). This clustering allowed us to label 10,000s of IRES sequences that were previously labeled largely based on species rather than IRES structure or not previously labeled at all. When comparing to the manual annotation there was substantial agreement (ARI = 0.78), but with the advantage of being done purely computationally.

Interestingly, within each type we saw subgroupings. We investigated what was driving the subclustering. Many subclusters resulted from Albratross missing a stem, but some of the divergent clusters resulted from *bona fide* newly predicted structures. One such cluster composed of 107 sequences including chicken Sicinivirus IRESes featuring a novel extended stem. We present Sicinivirus MW684796 as a specific example for follow-up experiments^66^.

The cluster retained hallmark Type II features including the GNRA-containing Domain I and the eIF4G-binding Domains J-K, with Domain J exhibiting high sequence and structure identity to EMCV (Figure 6C, D). However, the novel group harbored an extended AAG stem in Domain I not present in canonical Type II IRESes (Figure 6C–F).

The extended stem contains two conserved AAG motifs, with spacing from the RAAA stem to the first AAG matching that of EMCV, potentially enabling a similar contact with uS19 as recently reported^67^.

We performed DMS-MaPseq on the Sicinivirus MW684796 IRES, which experimentally validated that the IRES folds into the extended AAG stem conformation (Figure 6G–I). Indeed Domain I, containing the extended stem had particularly high chemical signal agreement to the predicted structure (Figure 6I).

We investigated the effects of the AAG stem extension on IRES activity. We cloned the Sicinivirus MW684796 IRES into our dual luciferase vector and transfected the construct into UMNSAH/DF1 chicken embryonic fibroblast cells. We also included series of mutants that shortened the extended stem. “Estem” replaced the extended AAG stem with the AAG stem from EMCV as a maximally conservative mutant. The *α* mutant shortened the stem to match the length and spacing of typical Type II architecture. The Δ mutant deleted the extended AAG stem entirely (Figure 6J).

The Sicinivirus MW684796 IRES exhibited low but measurable activity in this cell type as compared to the no IRES control (Figure 6K). Meanwhile, each of the extended AAG stem mutants were indistinguishable from the negative control, implying a feature in the extended stem is important for IRES activity in this subtype(Figure 6K).

### Dependency mapping learns generalizable structural patterns in a model size dependent manner

In natural language processing model size typically correlates with performance and we wanted to investigate if the same pattern applies to dependency analysis of RNA Language Models^68^. We followed a similar methodology as above on three version of RiNALMo: micro, mega, and giga (33M, 150M, and 650M parameters, respectively). For this experiment, we only trained on ∼38,000, Type I IRES sequences. Similar to natural language processing, when a Type I IRESes, the giga model predicted more stems correctly (Figure 7A).

**Figure 7:**
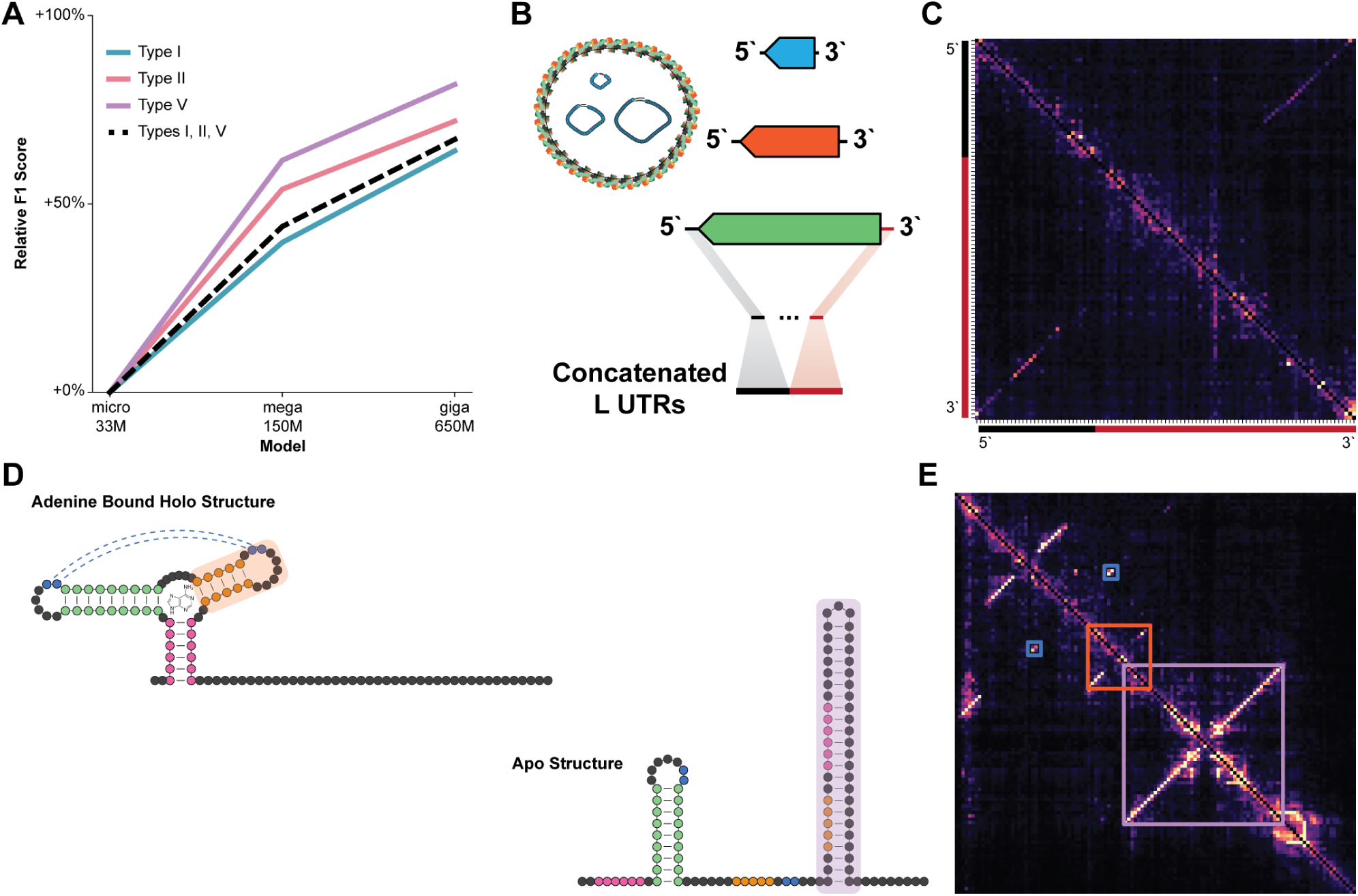
Generalizability of Albatross across RNA families. (**A**) Relative macro-F1 versus the micro model. One checkpoint was selected per model size by macro-F1 on Type I, II, and V. (**B**) Hantavirus virion schematic: the three genomic segments (L, M, S), coated by nucleocapsid protein (blue) and pseudo-circularized, packaged within the envelope glycoprotein lattice. Training input: hantavirus L segment UTRs, each with a 5’ (black) and 3’ (red) terminus, concatenated across strains for training. (**C**) Dependency map of the concatenated L UTRs from the 2026 outbreak isolate (PZ385161.1). The model correctly predicts pairing between the 5’ (black) and 3’ (red) termini, capturing pseudo-circularization, with no additional structure predicted. (**D**) Structures of the *B. subtilis* adenine riboswitch in the ligand-bound (holo) and unbound (apo) states. Shaded orange and shaded purple boxed stems correspond to the boxes in **E** . (**E**) Dependency map of the riboswitch. Boxed regions mark nucleotides with conflicting predicted pairing partners, consistent with the mutually exclusive holo and apo states in (**D**). The blue box marks the L2–L3 kissing-loop interaction, a tertiary contact common to both states, predicted alongside the state-specific pairs.

We tested the the Type I only models against Type II and Type V IRESes in our “ground truth” data. Model size correlated with improved F1 score, regardless of the IRES type (Figure 7A). This implies the larger models are able to better capture general rules that apply across IRES types.

Together these suggest a larger model could lead to more accurate and generalizable predictions of the secondary structure. Accordingly, this presents a strong argument for training larger models in future works.

### Continued pretraining of RNA language models enables structure prediction of diverse RNA families

In this work we showed that continued pretraining of an RNA language model on IRES sequences led to structure prediction with very high precision. However, we wanted to test if this approach generalizes to other RNA families.

In the midst of the 2026 hantavirus outbreak, turned our approach toward the novel strain^69^. Hantavirus genomes consist of three pseudo-circularized segments (L, M, S), each coated by the nucleoprotein for the majority of the life cycle (Figure 7B)^70^.

We continued pretraining a model on 5041 concatenated 5^′^+3^′^ UTRs drawn from S, M, and L segments across Orthohantavirus (Figure 7B)^38^. The resulting predictions on the novel strain correctly identified the long-range pseudo-circularization (Figure 7C). The model did not predict any additional structure, which is in line with the fact that hantaviruses RNA is coated by nucleoprotein for the majority of the life cycle and thus unable to for complex structure^70^.

Branching out beyond viral RNAs, we turned to bacterial riboswitches, RNAs that change structure in response to metabolites to control gene expression (Figure 7D)^71^. This family has the added complexity of inherently having two mutually exclusive functional states with conflicting base-pairing interactions.

We continued pretraining a model on 76,829 bacterial riboswitches spanning ten ligand classes, extending each aptamer 150 nt into its downstream expression platform^31,72,73^. The model recapitulated the canonical structure of the *B. subtilis* adenine riboswitch and predicted correct structures for other adenine riboswitches and for glycine riboswitches (Figure 7D, E, S11)^74,75^. Within a single map, it assigned individual nucleotides two mutually exclusive pairing partners that correspond to the apo and holo conformations. The model received no ligand, no structural label, and no indication that these RNAs adopt more than one state, and recovered the conformational switch from sequence alone. The same map also contained the L2–L3 kissing-loop interaction, the tertiary contact that holds P2 and P3 in parallel across the purine riboswitch family^76^ (Figure 7E).

Dependency maps are therefore not restricted to a single nested structure per sequence. They report the set of constraints an RNA is under, including cases where one nucleotide is held by two partners in two functional states, which positions this approach for discovery of RNAs whose function depends on conformational change rather than on a fixed fold.

Importantly, none of these structure features could be captured by base RiNALMo model. The Albatross strategy allows discovery of conserved structures across kingdoms of life using sequence-only continuous pretraining.

## DISCUSSION

This work aimed to study picornaviral IRESes as a comprehensive family rather than as individual RNAs. In doing so, we generated large-scale functisonal (96 IRESes across 6 cell types) and structural (∼75,000 dependency maps) atlases. These data are available in Supplementary Data S1 and at www.albatrossrna.org, respectively. Comparable cell-type-resolved activity measurements have been made for cellular 5^′^ and 3^′^ UTRs and used to design untranslated regions with tailored cell-type specificity^77^; our atlas provides the equivalent resource for full-length viral IRESes.

A recent study examining IRES context effects corroborates our activity rankings in overlapping cell types^48^. Across our atlas, Type V IRESes consistently showed strong activity in all cell types tested, and most IRESes exhibited significant tissue tropism. The strongest acting of these came from animal hosts, potentially because the human cells lack antiviral response mechanisms optimized by coevolution to suppress the activity.

The structure of the Type V eIF4G-binding domain offers a possible explanation for its activity. This domain is distinct from Type I and Type II in its overall architecture, but its apex carries a motif resembling the Type II eIF4G-contacting element alongside a GNRA tetraloop shared with domain V of the Type I Coxsackievirus B3 IRES^54,61^, the domain we compare against a crystal structure in Figure 3F–H. Combining elements of both other eIF4G-dependent classes may support more robust engagement of the initiation machinery, which would fit activity that holds across tissues rather than peaking in one. Type IV IRESes sit at the opposite end, ranking lowest in every cell type we profiled. These elements bind the 40S subunit directly, bypassing eIF4G and recruiting no ITAFs^78^, a comparatively minimal mechanism that may cap their output in this context. Distinguishing either explanation from alternatives requires mutating the individual motifs, as we did for the extended AAG stem in Sicinivirus MW684796, and measuring activity across far more IRESes than the 96 profiled here.

Using dependency mapping, we generated 75,000 IRES structural maps. Albatross was trained on sequence alone, given no knowledge of structure or thermodynamics, yet predicted base pairs with high precision, outperforming state-of-the-art methods by median precision and F1, with comparable recall. Notably, the Infernal combines curated covariation with folding grammar^79,80^, itself trained on thousands of known RNA structures, whereas Albatross received no structural information of any kind. This advantage likely stems from covariation analysis yielding a single consensus structure per family, which is then mapped onto the query sequence. Dependency mapping instead predicts a structure for each individual sequence, capturing features that a family consensus cannot represent. Divergence remains a constraint on covariation approaches, since a family must first be recognized and populated with enough homologs to model as illustrated by the Gallivirus IRES. Albatross requires no family assignment or homolog set, making it applicable to divergent sequences that covariation methods cannot reach.

Crucially, through multiple experiments, we demonstrate that the Albatross dependency mapping approach learns generalized patterns that it then can apply to divergent but related contexts (Figure 4D–G, 5, 7A). Furthermore, this approach has high precision and preferentially predicts structures using sequence context (Figure 3A–C, 5, Table 2). These properties are particularly useful in comparison to pure thermodynamic models as the Albatross predicted stems are more likely to have functional significance.

A fraction of “false positive” pairs are due to the model extending a predicted stem through what the canonical structure marks as a bulge(Figure S11) because the dependency map shows a clear coupling between the two bases that would otherwise sit unpaired across from each other (*e.g* A • C). A thermodynamic model will not pair these, which suggests that Albatross reports on the spatial coupling between nucleotides proximal in three-dimensional space.

The Albatross approach toward RNA structure prediction has the additional strength of depending only on sequence data. As compared to high-quality RNA structure data, sequences are orders of magnitude more readily available or obtainable. For example, while only a handful of IRES structures have been experimentally resolved, Albatross generated *>*75,000, enabling comparison at scale.

In analyzing the bulk data, using a new structure clustering method, we discover a previously unreported subclass of IRESes with an extended AAG stem. We experimentally confirm the extension and find it is enhances IRES activity in the context of this subclass (Figure 6G–K).

Overall, we provide a family-scale analysis of IRESes through multiple high-throughput avenues. Our functional data can be applied to biotechnology and infectious disease research investigating picornaviral tropism. Meanwhile, we establish continued pretraining of RNA language models with dependency mapping as a viable approach for high-throughput structural analysis of diverse RNA families (Figure 7B–E).

Our work demonstrates that information on functional RNA structure is contained and can be learned from sequences alone. This likely comes from selective pressure that gave raise to the particular arrangement of nucleotides, which survived millions of years of evolution. The approach we develop does not recognize all stable structures but very specific conserved ones, which makes this analysis broadly applicable for discovery of functional RNA structures and structure switches across multiple kingdoms of life.

### Limitations of the study

This study focused picornaviral IRESes and did not include IRESes or similar non-canonical translation elements from other sources. Type III IRESes are relatively homogeneous in sequence and can be more easily mapped to the canonical structure. Nonetheless our ground truth structure data set lacked Type IIIs so we do not test model performance on this type. Albatross’s high precision is offset slightly by a lower recall. Though this is a useful feature for constricting the search space for functional structures, it also means some substructures are likely missed in any given prediction. The Sicinivirus MW684796 IRES was tested in an easily accessible and transfectable gallinaceous cell type where it exhibited weak activity. Future follow ups on the mechanisms employed by the extended stem will need to evaluate other cell types to better assess the IRES function. Finally, we use a generous definition of IRES throughout for what is more precisely a “picornaviral 5^′^ UTR”.

## RESOURCE AVAILABILITY

### Lead contact

Requests for further information and resources should be directed to and will be fulfilled by the lead contact, Silvi Rouskin.

### Materials availability

Plasmids generated in this study are available upon request from the lead contact.

### Data and code availability

- DMS-MaPseq and polysome profiling sequencing data have been deposited at GEO and are publicly available as of the date of publication under accession number GEO: GSE34725 (DMS-MaPseq) and GSE347508 (poysome profiling). Albatross dependency maps for approximately 75,000 IRESes are available at https://albatrossrna.org, and DMS-MaPseq derived structure models are available at https://www.rnandria.org.
- All original code used for training is available at https://github.com/rouskinlab/albatros
- Albatross model weights, and the sequence datasets are archived at Zenodo: https://doi.org/10.5281/zenodo.22920917.
- Any additional information required to reanalyze the data reported in this paper is available from the lead contact upon request.

## ACKNOWLEDGMENTS

We thank Julien Gagneur for providing insight and suggestions for applying dependency mapping to a new context. We further thank Mélissa Léger-Abraham for allowing us to use the gradient station in her lab. Additionally, we thank Olivier Pourquié for providing the UMNSAH/DF1 cells.

A. S. was partially supported by the National Institute of Allergy and Infectious Diseases Training Grant NIHT21AI007245-40. A. S. and P. B. were partially supported by the Giovanni Armenise Foundation. This research was supported by the the Giovanni Armenise Foundation, Massachusetts Consortium on Pathogen Readiness, Massachusetts Life Sciences Center, and the Advanced Research Projects Agency for Health.

## AUTHOR CONTRIBUTIONS

Conceptualization: A.S., R.A.W, N.B., S.R. Data Curation: A.S., P.B., G.Y., A.I. Formal Analysis: A.S., P.B., A.I. Funding Acquisition: A.S., S.R. Investigation: A.S., P.B., G.Y., A.I., S.R. Methodology: A.S., P.B., J.R. R.A.W., S.R. Project Administration: A.S., S.R. Resources: A.S., P.B., G.Y., R.A.W Software: A.S., P.B. Supervision: S.R. Validation: A.S., P.B., G.Y. Visualization: A.S., P.B. G.Y. Writing – Original Draft: A.S., P.B., G.Y., S.R. Writing – Review & Editing: A.I., J.R., R.A.W., N.B.

## DECLARATION OF GENERATIVE AI AND AI-ASSISTED TECH-NOLOGIES

The authors did not use generative AI for writing initial draft but AI tools were used for grammar, flow editing, condensing word count, and catching errors within the text. Tools including Claude and Codex, were also used for assistance in coding. The authors reviewed the code to ensure that it was accurate.

## SUPPLEMENTAL INFORMATION INDEX

Figures S1-S11 and their legends in a PDF

Data S1. Raw data for polysome traces, qPCR, and library composition. Processed data for ribosomal loading and DMS-MaPseq. Sequence property summaries of IRESes and A71 synthetic replacement experiments.

## STAR METHODS

### Experimental model and study participant details

#### Cell lines

All cell lines were grown in DMEM (ThermoFisher Scientific #10564011) with 10% FBS (ThermoFisher Scientific #A5209402) supplementation at 37°C with 5% CO_2_. UMNSAH/DF1 cells were grown at 39°C with 5% CO_2_.

The following cell lines were used: HEK293T, Female; U2OS, Female; SH-SY5Y, Female; HCT-8, Male; Huh7, Male; N2A (Neuro-2a), Male, (*Mus musculus*, A/J); UMNSAH/DF1, Unspecified, (*Gallus gallus*, East Lansing Line 0). Cells were purchased commercially, except UMNSAH/DF1. The UMNSAH/DF1 cell identity was confirmed using RNAseq alignment to *Gallus gallus* genome.

### Method details

#### Plasmids

All PCR reactions were performed using CloneAmp HiFi PCR Premix (Takara Cat. #639298). Isothermal Assembly was performed using NEBuilder HiFi DNA Assembly Master Mix (NEB E2621L). Constructs were chemically transformed into NEB Stable Competent E. coli (NEB C3040I) or electroporated into NEB 10-beta Competent E. coli (NEB C3019I).

#### RT-qPCR

cDNA was synthesized using LunaScript RT SuperMix (NEB M3010L). RT-qPCR was performed using either either Luna Universal Probe qPCR Master Mix (NEB M3004L) for probe-based or Luna Universal qPCR Master Mix (NEB M3003) for SYBR-based RT-qPCR as noted throughout.

#### Next-generation sequencing

Next-generation sequencing library preparation was performed using either NEBNext UltraExpress RNA Library Prep Kit (NEB E3330L) or NEBNext UltraExpress DNA Library Prep Kit (NEB E3325L) as appropriate. Next-generation sequencing was performed on a Illumina NextSeq using NextSeq 1000/2000 P1 XLEAP-SBS Reagent Kit (300 Cycles) (Illumina 20100982).

#### circRNA Plasmid Library

IRES sequences were obtained from Twist Bioscience in 96-well plate format. Each IRES included overhangs to clone into the circRNA generation plasmid provided by the Mass General Hospital RNA Therapeutics Core. The donor circRNA plasmid (p10-Natural-RTC) was amplified using and primers oAS092 and oAS093. IRES and donor plasmid were assembled together using isothermal assembly and transformed into NEB 10-beta Competent E. coli (NEB C3019I).

The distribution of IRESes within the plasmid library was confirmed by Next-Generation Sequencing a PCR amplicon of the barcode region associated with each IRES (Primers oAS052, oAS077). These data recapitulated the results of sequencing randomly fragmented library by NEBNext dsDNA Fragmentase (NEB M0348S).

#### circRNA library generation and analysis

circRNAs were generated by the Mass General Hospital RNA Therapeutics Core. Distribution within the circRNA library was measured by generating cDNA then sequencing a PCR amplicon of the barcode region. This was validated by sequencing a randomly fragmented library . Circularization was confirmed by probe-based RT-qPCR using primers spanning the splice junction.

#### Polysome profiling

Polysome gradient buffer (PGB) consisted of 20mM Tris-HCl pH 7.5 (Rockland #MB-002), 100mM KCl (Invitrogen AM9640G), 10mM MgCl_2_ (Thermo Scientific J62411.AD). 10% and 50% sucrose solutions in PGB were set up. 13.2 mL Open-Top Thinwall Ultra-Clear Ultracentrifuge Tubes (Beckman Coulter 344059) were loaded with the 10% sucrose solution up to the level indicated by the BioComp Marker Block (BioComp Instruments 105-914A-R). Using a syringe attached to cannula the 50% sucrose solution was loaded to the bottom of the tube until the total solution was level with the BioComp Gradient Station MagnaBase Tube Holder (BioComp Instruments 105-914A-R). The tubes were capped and the SW41 Rotor, Long Sucrose 10-50% L11 protocol was run on the BioComp Gradient Station (BioComp Instruments 153).

10 cm plates containing transfected cells were treated with 100 *µ*g/mL cycloheximide (Sigma-Aldrich C4859-1ML) for 10 minutes at 37°C. Lysis buffer was made with 1 mL PGB, 500 *µ*g cycloheximide (Sigma-Aldrich C4859-1ML), 5 *µ*L NP-40 (BioBasic NDB0385-100ml). 500 *µ*L of lysis buffer was added to each 10 cm plate. Cells were scraped and transfered in to 1.7 mL tubes on ice for 10 minutes. The lysis was spun down at 4°C at 11,200 rcf for 5 minutes and supernatant collected into clean tubes.

150 *µ*L of the lysate was loaded on to each sucrose gradient. The gradient was spun in a SW 41 Ti Rotor (Beckman Coulter 331362) in an L100XP ultracentrifuge (Beckman Coulter 392052) at 36,000 rpm (221,356 rcf) for 1 hour 20 minutes.

Sucrose gradients were fractionated using the BioComp Gradient Station (BioComp Instruments 153) collecting fractions that corresponded to polysome peaks.

RNA was extracted using 1 mL TRIzol Reagent (ThermoFisher Scientific 15596026) then following manufacturer protocol. cDNA was synthesized as above and barcodes were amplified via PCR reactions using CloneAmp HiFi PCR Premix (Takara Cat. #639298) for 25 cycles. Amplicons were sequenced by next-generation sequencing as above. RNA from the lysis was similarly processed.

Barcodes counts were normalized to the total reads then normalized to the fractional representation in the lysis.

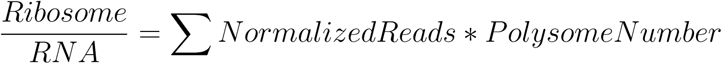

#### Luciferase assays

Cells in a 96 well plate were transfected with 10 ng plasmid DNA via PEI protocol (MedChem Express HY-K2014). Twenty four hours later, media was removed and 80 uL of 1x Passive Lysis Buffer (Promega Cat.# E1910) was added to the cells. Luciferase assays were performed using the Nano-Glo Dual-Luciferase Reporter Assay System (Promega Cat.# N1610) according to manufacturer protocol.

#### DMS-MaPseq

For *in vitro* DMS-MaPseq, RNA was suspended to 0.5 *µ*g/*µ*L in 10 L of H2O, incubated at 95°C for 1 min, immediately mixed with 87 *µ*L of 37°C preheated refolding buffer (0.4M Sodium cacodylate (Electron Microscopy Sciences #11655), 6 mM MgCl2 (Thermo Scientific J62411.AD)). The reaction was incubated at 37°C for 30 min. 3% DMS (Sigma-Aldrich #D186309-100ML) was added to each reaction and incubated at 37°C for 5 min before being quenched with 0.6 volumes *β*-Mercapethanol (Thermo Scientific A15890.0B). RNA was isolated using Monarch Spin RNA Cleanup Kit (NEB T2030L). Next-generation sequencing library preparation was performed using either NEBNext UltraExpress RNA Library Prep Kit (NEB E3330L) but replacing the reverse transcriptase with Induro Reverse Transcriptase (NEB M0681L).

For *in cellulo* DMS-MaPseq, 5 *µ*g RNA was transfected into HEK293T, N2As, or HCT-8s in 6-well plates. After 6 hours, cells were treated with 3% DMS (Sigma-Aldrich #D186309-100ML) for 5 minutes at 37°C. The reaction was quenched with 10 mL 30% *β*-Mercapethanol (Thermo Scientific A15890.0B) in DPBS (ThermoFisher Scientific #14190144). RNA was isolated using TRIzol extraction and libraries were prepared as described above.

DMS-MaPseq sequencing data was processed using SEISMIC v24.3 (https://github.com/rouskinlab/seismic-rna) to map DMS reactivity, cluster alternative structures, and evaluate secondary structure models^45^.

#### RNA Language Model

We used as a base model RiNALMo giga a 650 Million parameter RNA Language Model.

RiNALMo’s architecture was not modified. It is composed of 33 transformer blocks, 1280 embedding dim, 20 attention heads. It uses Flash attention. The attention dropout and residual dropout are both set at 0.1. RiNALMo was originally trained on 36M non-coding RNAs, none of which are IRESes.

The tokenization was the same as in the original RiNALMo. Each nucleotide is a single token. After collecting the IRESes two transformations were applied: First, the sequence would be uppsercased. Second, uracils were converted to thymines. These transformations were applied to stay in line with RiNALMo’s pretrainining. Therefore, the Albatross vocaulary is made of 4 vocabulary tokens: A, C, G, T; and four special tokens: [CLS], [EOS], [PAD], [MASK], and [UNK].

#### Training data & Preprocessing

Initially we collected the 5^′^ UTRs of every complete picornaviral genome in the NCBI Virus Database^38^. We then iteratively used BLAST to find the top 500 aligning sequences until we had a total of about 200,000 unique sequences^39^. To make the 500,000 sequence dataset, we used cd-hit to cluster sequences by 70% or greater sequence similarity^60^. We then ran an iterative BLAST on the least represented sequences until we had a total of about 500,000 unique sequences. The 50,000 sequence dataset was generated by using cd-hit to cluster the 500,000 dataset by 70% or greater sequence similarity^60^. The representative sequence of each cluster was added to the dataset, then the most divergent sequence of each cluster was added one at a time until the set included about 50,000 sequences.

Sequences collected containing any IUPAC ambiguity code were filtered out. Sequences having an exact match to another sequence were also filtered out, in order to avoid overrepresention.

The three datasets used in figure 3 are respectively composed of 193,643, 509,944, and 52,129 unique sequences. Three datasets were used throughout this paper; they are available on https://github.com/rouskinlab/.

#### Model Training

We downloaded the official released weights from RiNALMo and continued pretraining of all 650M parameters. We did not use any LoRA, adapters, or layer freezing. Training used sequence input only. No structural labels, base pairing rules or thermodynamic priors were given to the model or trained on.

The Objective is the same one as used in RiNALMo: a BERT-style masked language modeling at 15% mask rate. This mask would then have the following distribution: 80% [MASK] / 10% random nucleotide / 10% unchanged. Loss was computed on an independent 15% of nucleotides.

The optimizer used was AdamW with parameters: *β* = (0.9, 0.98), *ε* = 10^−6^, learning rate was 5 × 10^−6^ and weight decay was 0.01^81^.

A linear warmup was set over 10,000 steps from 0.01 · lr to lr. There is no decay phase, the learning rate is constant after warmup.

The training parameters were: Batch size = 8, Max sequence length = 2048, Gradient clipping 1.0, 16-mixed precision, Random seed 42, Up to 10 epochs.

Every experiment was run on a single L40 GPU. The 50k run completed in approximately twelve hours.

The weights released with this paper are from the 50k dataset, epoch 5, step 39,096. All dependency maps used in this paper and available on www.albatrossrna.org were produced from this checkpoint.

#### Dependency Map Generation

The Methodology used to get the dependency map is the same as described in da Silva et al.^35^.

For a sequence of length *N*, we run 3*N* + 1 forward passes (1 for the reference sequence and every single-point substitution to each of the 3 alternative bases). For each of these forward passes, we extract softmax probability over {*A, C, G, U* } at every position *j*. For mutation at position *i* to base *m* and target base *t*, compute the log-odds shift

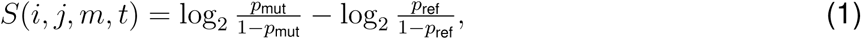

where *p*_mut_ = *P* (*x_j_* = *t* | *i* → *m*) and *p*_ref_ = *P* (*x_j_* = *t* | wild type) are the model’s softmax probabilities at position *j* for target base *t*, under the mutant and reference sequences respectively. The dependency map entry is *M* [*i, j*] = max*_m,t_* |*S*(*i, j, m, t*)|, with *M* [*i, i*] = 0. Each entry is directed: *M* [*i, j*] measures the effect of substituting position *i* on the prediction at *j*. Maps are not symmetrized at this stage.

#### Binary Structure Filtering

Binary filtering and structure prediction consisted of the following algorithm:

**Algorithm 1: Binary structure filtering from a dependency map.**

**Input:** dependency map *M* ∈ R*^N^*^×^*^N^* ; thresholds *τ, γ, α*.

**Output:** binary contact matrix *B* ∈ {0, 1}*^N^*^×*N*^ .

1. Normalize: *M* ^′^ ← clip(*M/*20, 0, 1).
2. Symmetrize: *M* ^′^ ← max(*M* ^′^*, M* ^′⊤^).
3. Threshold: *M* ^′^[*i, j*] ← 0 where *M* ^′^[*i, j*] *< τ* .
4. Diagonal-band exclusion: *M* ^′^[*i, j*] ← 0 where |*i* − *j*| ≤ *γ*.
5. Anti-diagonal connectivity: along each anti-diagonal {(*i, j*) : *i* + *j* = *s*}, retain only entries belonging to a contiguous run of length ≥ *α*.
6. Maximum-weight matching: build an undirected graph with edge weights *M* ^′^[*i, j*] and solve *B* ← arg max_M_ Σ_(_*_i,j_*_)∈M_ *M* ^′^[*i, j*] via Edmonds’ Blossom algorithm. This enforces one-to-one pairing and allows non-canonical pairs.
7. Pseudoknot removal: sort accepted pairs by span |*j* − *i*| in decreasing order, then iterate; retain a pair only if it does not cross any already-retained pair.

As Blossom matching can return crossing pairs, we removed pseudoknots to yield strictly nested structures representable in dot-bracket notation.

The filtering applied on the dependency map only encodes geometric priors and no biological priors. It is a significant design choice as it makes our prediction entirely dependent on the LLM log odds shifts. Therefore, the model has no idea of base pairing rules or thermodynamic priors. The dependency maps are raw and not symmetrized.

Each of the parameter was designed to encode for a different prior: *α* forces stem to have a minimum length. This prevents the filter to select isolated base pairs that also can be noise from the dependency map. *γ* excludes the base pairs immediately next to the diagonal, as these pairs cannot form (Triangle inequality). *τ* is the most significant parameter as it is the threshold to filter out weak signal.

#### Filter Parameter Selection (5-fold cross-validation)

In order to select the best set of parameters, we ran a 5-fold cross validation with a grid search on the 95 DMS-MaPseq-constrained IRES structures.

The 5-fold cross validation ran under the following settings: The 95 reference sequences were split using KFold with *K* = 5, shuffle=True, and random_state=42. For each fold, true positives, false positives, and false negatives were aggregated across the held-out sequences, and a micro-F1 score was computed. We then reported the mean and standard deviation across folds. The grid search covered *α* ∈ {3*, . . . ,* 8}, *τ* ∈ [0, 0.20], and *γ* ∈ {3, 4, 5}, with micro-F1 used as the metric to evaluate each parameter combination.

#### Selected parameters and performance (50k, epoch 5)

The selected parameters were *α* = 3, *τ* = 0.11, and *γ* = 4. Using these fixed parameters, five-fold cross-validation gave a micro-F1 of 0.57 ± 0.03, micro-precision of 0.79 ± 0.02, and micro-recall of 0.44 ± 0.03. Per-IRES scores for this checkpoint are reported in Table 1: F1 0.54 ± 0.21 (median 0.60), precision 0.73 ± 0.22 (median 0.80), and recall 0.44 ± 0.19 (median 0.48).

#### Structure comparison

We compared the predicted structures and reference structures as binary pairing matrices over the upper triangle. We used the standard definition of Precision, recall, and F1 such that *P* = TP*/*(TP + FP), *R* = TP*/*(TP + FN), and F1 = 2*PR/*(*P* + *R*).

#### Infernal Structure Analysis

We used Infernal 1.1.5. For each sequence we used cmsearch against the Rfam.cm database. cmfetch pulled the relevant model for each sequence and cmalign with --mxsize 8192 aligned each sequence to its corresponding model, producing the structural alignment used as input to R-scape. R-scape 2.6.4 was run on each alignment with CaCoFold enabled to generate the predicted structure. Unless otherwise noted all commands were run with default settings. The consensus structure was matched to the query sequence for F1, precision, and recall calculations. When Infernal returned more than one possible model, the best-scoring model by F1 was used, so the reported Infernal performance represents an upper bound. Because Rfam covariance models are built from curated family alignments, cmalign places each query sequence within that family’s alignment, so the resulting CaCoFold structure closely tracks the Rfam consensus structure for that family.

#### eFold structure prediction

We used eFold released model weights for each of the 95 sequences to predict their secondary structure^23^. The default post-processing and code of eFold was used. All the sequences were converted to the RNA alphabet, no DMS reactivities were fed to the model. The evaluation of precision, recall, and F1 were done using the same methodology as prior.

#### Unconstrained thermodynamic structure prediction

We used Fold from RNAstructure version 6.4 to generate thermodynamics predictions of the 95 sequences^21^. Each sequence was folded at 310.15 K (37 ^◦^C) using the minimum-free-energy structure. The rest of the parameters were default parameters. No DMS reactivities or base-pair constraints were given to the RNAstructure. The predicted structures of each sequence were converted to binary pairing matrix and evaluated with the same methodology described above.

#### Albatross-Constrained Thermodynamic Structure Prediction

The base pairs generated from Albatross with standard parameters were given as hard constraints to RNAstructure Fold version 6.4^21^. The predictions given to the thermodynamic model were restricted to AU, GC, and GU pairs. The sequences were then folded at 310.15 K (37 ^◦^C) using the constrained minimum-free-energy structure without DMS reactivities.

#### Family-Wide Structure Maps

Across the full set of binary maps, we observed a mean of 71.5 base pairs per map, a median of 78, and a maximum of 171. After substring-containment deduplication of the 500k pool, 75,229 root sequences were retained and used for the family-wide structural analyses. These structures can be found on https://albatrossrna.org.

#### A71 model for synthetic sequence testing

For the data in Figure 5, the model was trained as above on 8,553 A71 sequences, which represented the with 70% similarity using cd-hit^60^.

Tested synthetic sequences are available in the Supplementary Data and were processed as described in dependency map generation above.

#### Structure alignment and clustering

Up to 10 sequences were selected from each cd-hit 70% similarity cluster. Binary maps were used as *n x n* adjacency matrices. Each antidiagonal was projected into the corresponding column of a *n x* (2*n* − 1) matrix and padded with zeros. Columns with no pairs were dropped.

The resulting compressed structure matrices were aligned using a modified Needleman-Wuncsh Algorithm with a gap penalty of -1^82^. Aligned columns *a, b* were scored as 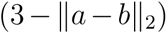.

For structure clustering, the final modified Needleman-Wuncsh similarity score between structures *i, j*, *S_i,j_ ɛS* was converted *d* 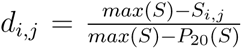. *d* was used as the distance for hierarchical clustering using scipy.cluster.hierarchy.linkage with method = “average”.

#### Generalization

In order to test generalization (Figure 7A), we continued pretraining the three model sizes from RiNALMo micro (33M), mega (150M), and giga (650M) on type I IRES sequences. The dataset was made of 38,223 sequences. These sequences were selected from the 75,229 unique set of IRESes. No Type II, III, IV, nor V IRESes were part of the training. The training settings were the same as described above.

The dependency maps were also produced using the same methodology. The maps were produced every 3,125 steps for the 85 Type I, II, and V sequences for which we have *in cellulo* DMS-MaPseq ground-truth. The dataset is composed of: 56 Type I, 16 Type II, 13 Type V. The parameters used to turn the dependency map into a binary map were the same as in the previous section: *α* = 3, *τ* = 0.11, and *γ* = 4.

For Figure 7A, one checkpoint was selected per model size using the macro-F1 on the Type I, II, and V sequences (85 sequences). The checkpoints were: micro 62,500; mega 50,000; giga 53,125. The panel shows the relative micro-F1 versus the micro model.

#### Hantavirus

We collected 15,375 Orthohantavirus nucleotide records from NCBI. As RiNALMo was only trained on non-coding sequences, we had to filter out on coding regions. Only 14,884 sequences had annotated CDS. For each unique sequence, we labeled the left of the CDS: 3^′^UTR and the right of the CDS: 5^′^UTR. We then, for each of this independent segment, concatenate them. Of the 14,884 CDS-annotated records, 9,843 had no sequence flanking the CDS, so the concatenated UTR was empty and those records were removed. After this filtering, we had: 5,041 sequences. In case of ambiguity code being present in the sequence, the letter would be replace by an N.

We continued pretraining the RiNALMo giga model as above but with a learning rate of 5 × 10^−6^, a batch size of 8, and 60,000 training steps.

The dependency maps were generated for the S, M, and L segments of ANDV/Switzerland/Hu-3337/2026 using the same methodology. For each segment, the genomic 3^′^ and 5^′^ UTRs were concatenated in the same manner as the training sequences.

#### Dependency mapping of bacterial riboswitch

We continued pretraining RiNALMo on 76,829 bacterial riboswitch sequences from 10 ligand classes taken from the RFAM full-region alignments for RF00050, RF00059, RF00162, RF00167, RF00168, RF00174, RF00504, RF00522, RF01051 and RF01734^72^. Since Rfam covariance models contain only the aptamer domain and not the corresponding downstream expression platform which contains the switch, we extended each hit with 150nt to the 3’ of the aptamer from the corresponding full sequence in NCBI^73^. The *b. subtilis* riboswitch dependency map was generated as described in dependency map generation above using this model.

### Quantification and statistical analysis

Statistical analysis was performed using python 3.11.5, numpy 2.2.6, scikit-posthocs 0.11.4, and scipy 1.17.1.

Statistical methods are noted in relevant figure legends. All error bars represent standard deviation from the mean.

### Additional resources

Albatross Dependency Map Database: https://albatrossrna.org

DMS-MaPseq Data: www.rnandria.org

## SUPPLEMENT FIGURES TITLES AND LEGENDS

**Figure 1:**
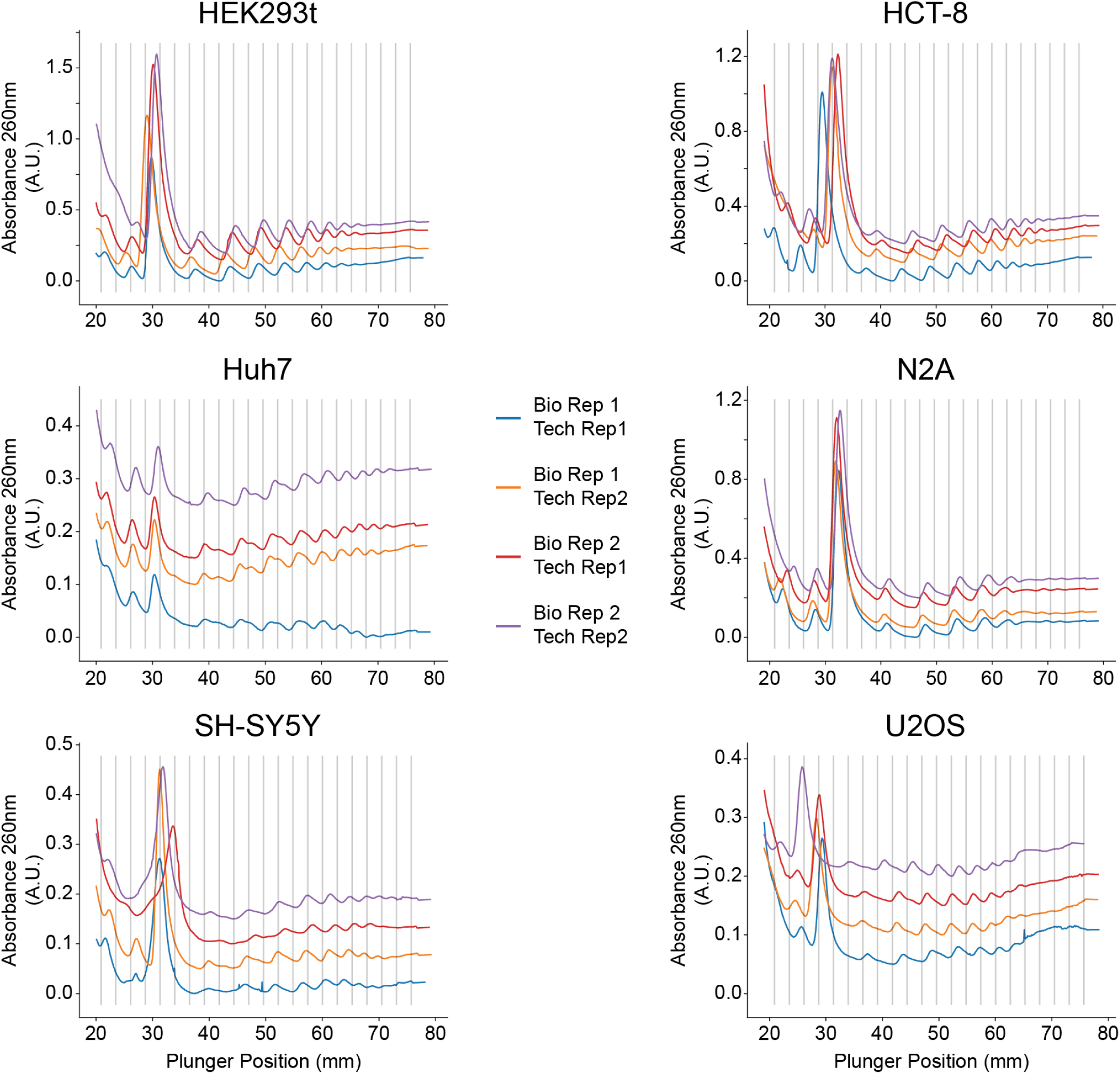
Polysome fractionation absorbance at 260nm. Trace of 260 nm absorbance during polysome fractionation. For each sample, absorbance was shifted equally at each point to set minimum absorbance to 0 A.U. Replicates were similarly shifted upwards for clarity. Vertical lines delineate fractions.

**Figure 2:**
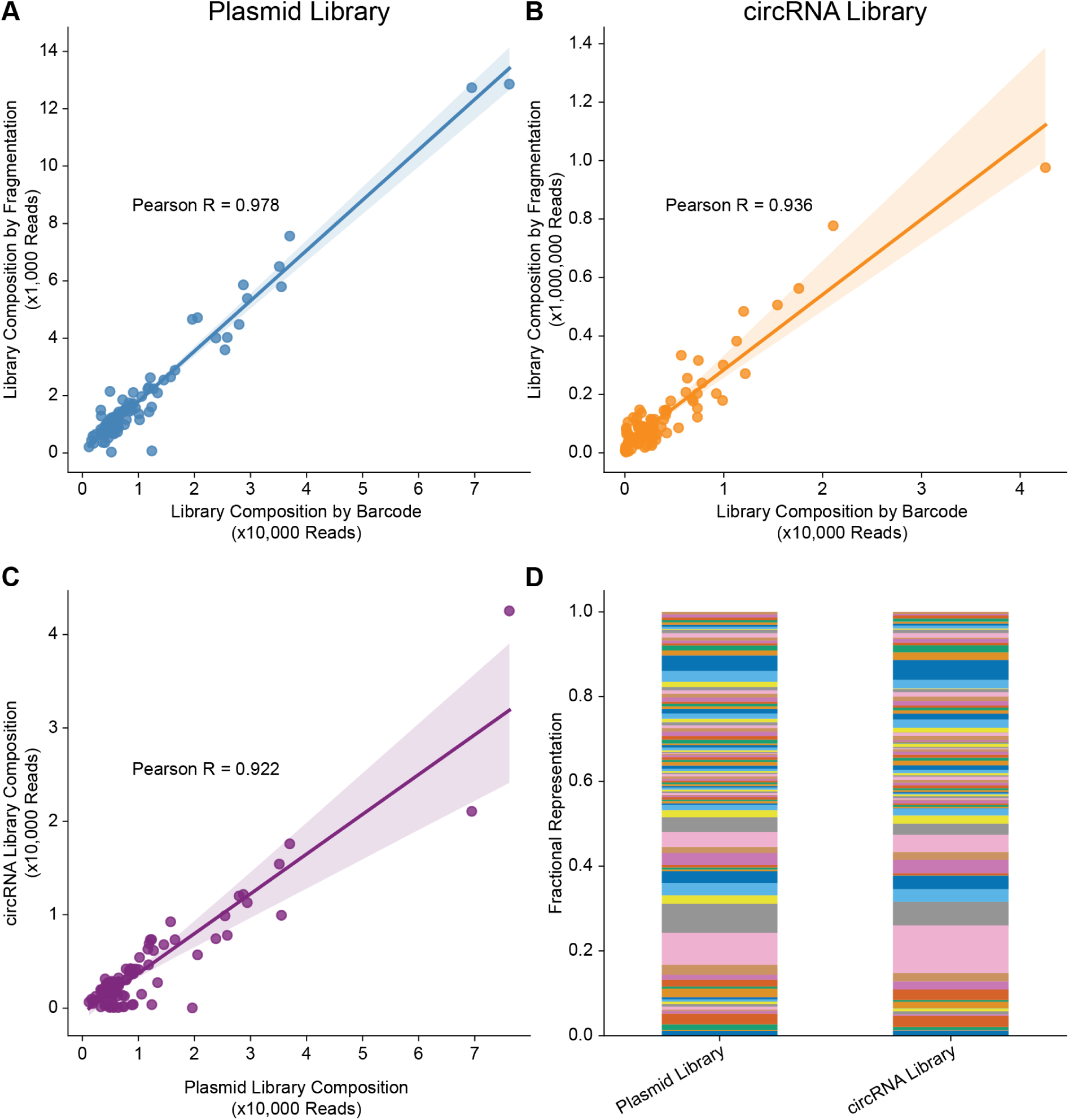
IRES representation is consistent across library preparation and sequencing methods. (A) Correlation between barcode sequencing and random fragmentation sequencing for the plasmid library (Pearson R = 0.978). (B) Same as (A), but for the circRNA library (Pearson R = 0.936). (C) Correlation between plasmid and circRNA library composition measured by barcode sequencing (Pearson R = 0.922). Each point represents one IRES. (D) Fractional representation of individual IRESes in the plasmid and circRNA libraries. Each colored segment corresponds to a single IRES. Shading is consistent between libraries. Lines indicate linear regression fits with 95% confidence intervals.

**Figure 3:**
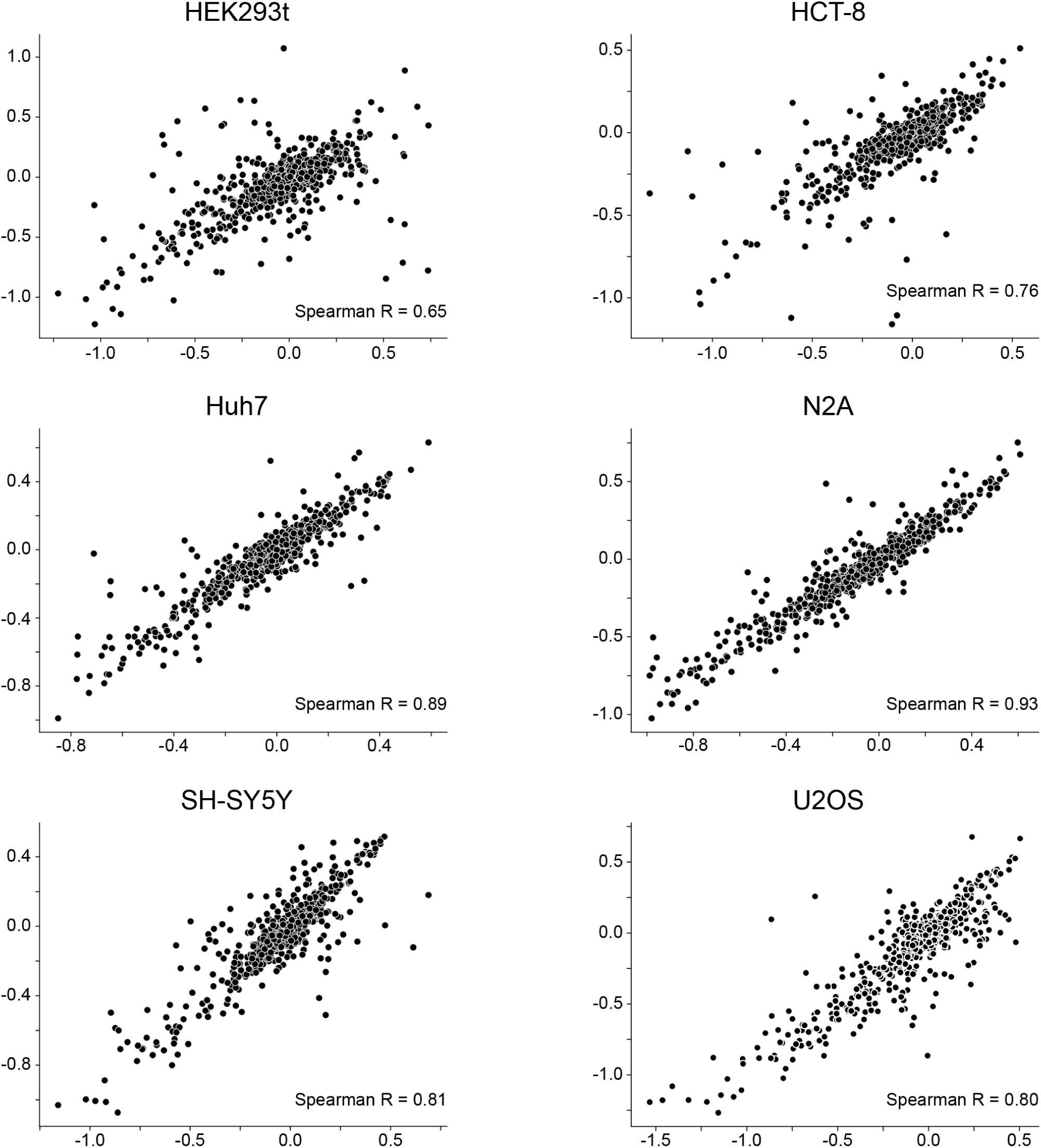
Polysome profiling barcode abundances are reproducible across replicates. Scatter plots of barcode abundances between biological replicates for each cell type. Spearman correlation coefficients are indicated: HEK293T (R = 0.65), HCT-8 (R = 0.76), Huh7 (R = 0.89), N2A (R = 0.93), SH-SY5Y (R = 0.81), and U2OS (R = 0.80). Each point represents one IRES barcode in a given polysome fraction.

**Figure 4:**
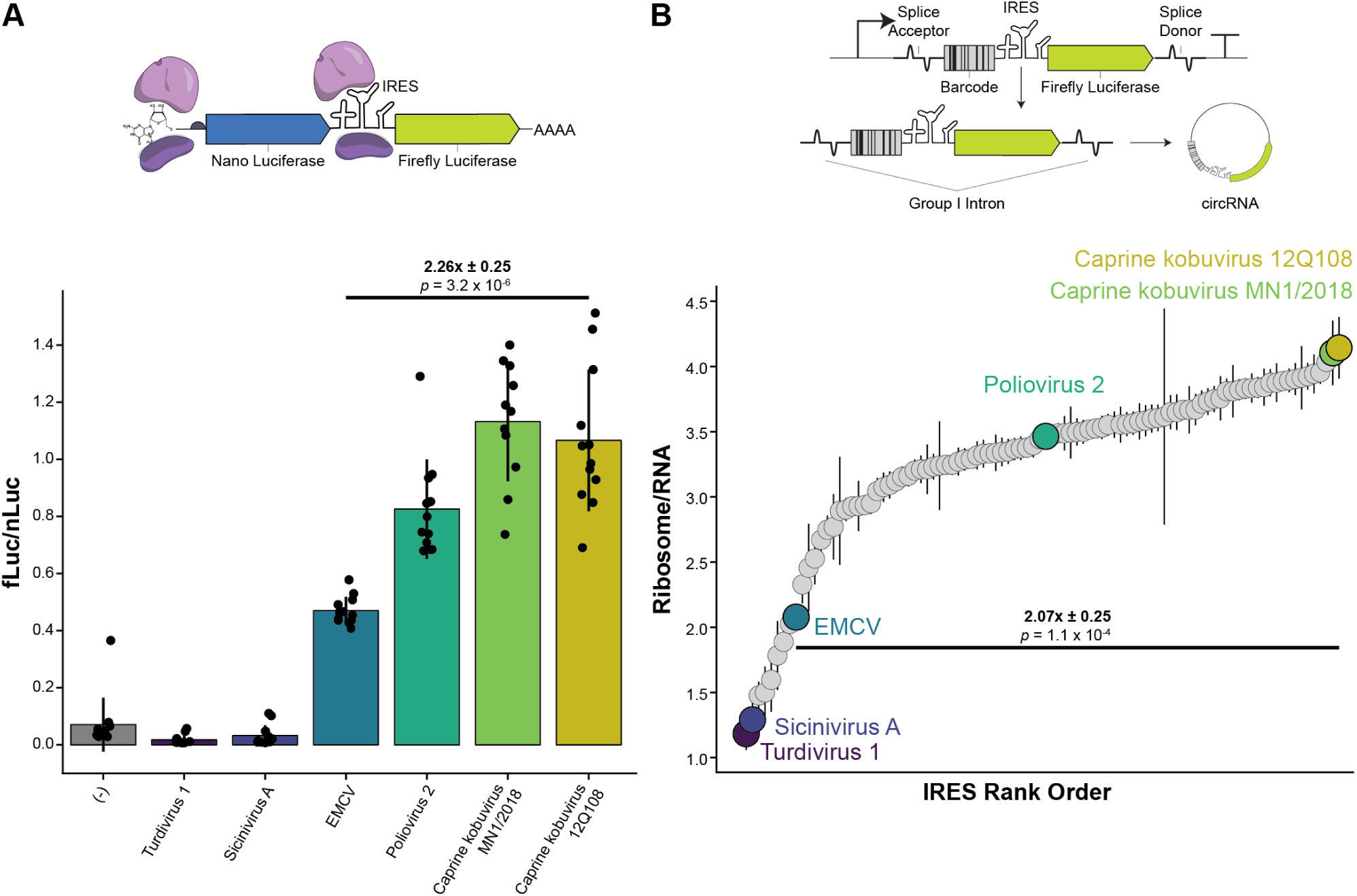
Dual luciferase reporter assay validates polysome profiling rank order. (A) Bicistronic dual luciferase reporter assay schematic (left) and firefly to nano luciferase ratios (fLuc/nLuc) for six selected IRESes: the top two (Caprine kobuvirus 12Q108, Caprine kobuvirus MN1/2018), bottom two (Turdivirus 1, Sicinivirus A), median (Poliovirus 2), and EMCV. Caprine kobuvirus 12Q108 exhibits 2.26 *±* 0.25 fold activity relative to EMCV (*p* = 3.2 *×* 10*^−^*^5^). Data points represent individual replicates; bars indicate the mean. (B) CircRNA construct schematic (top) and ribosomes/RNA from polysome profiling for all 96 IRESes ranked by activity, with the same six IRESes highlighted. Caprine kobuvirus 12Q108 exhibits 2.07 *±* 0.25 fold activity relative to EMCV (*p* = 1.1 *×* 10*^−^*^3^). Rank order and relative expression are concordant between the two assays. All data from HEK293T cells.

**Figure 5:**
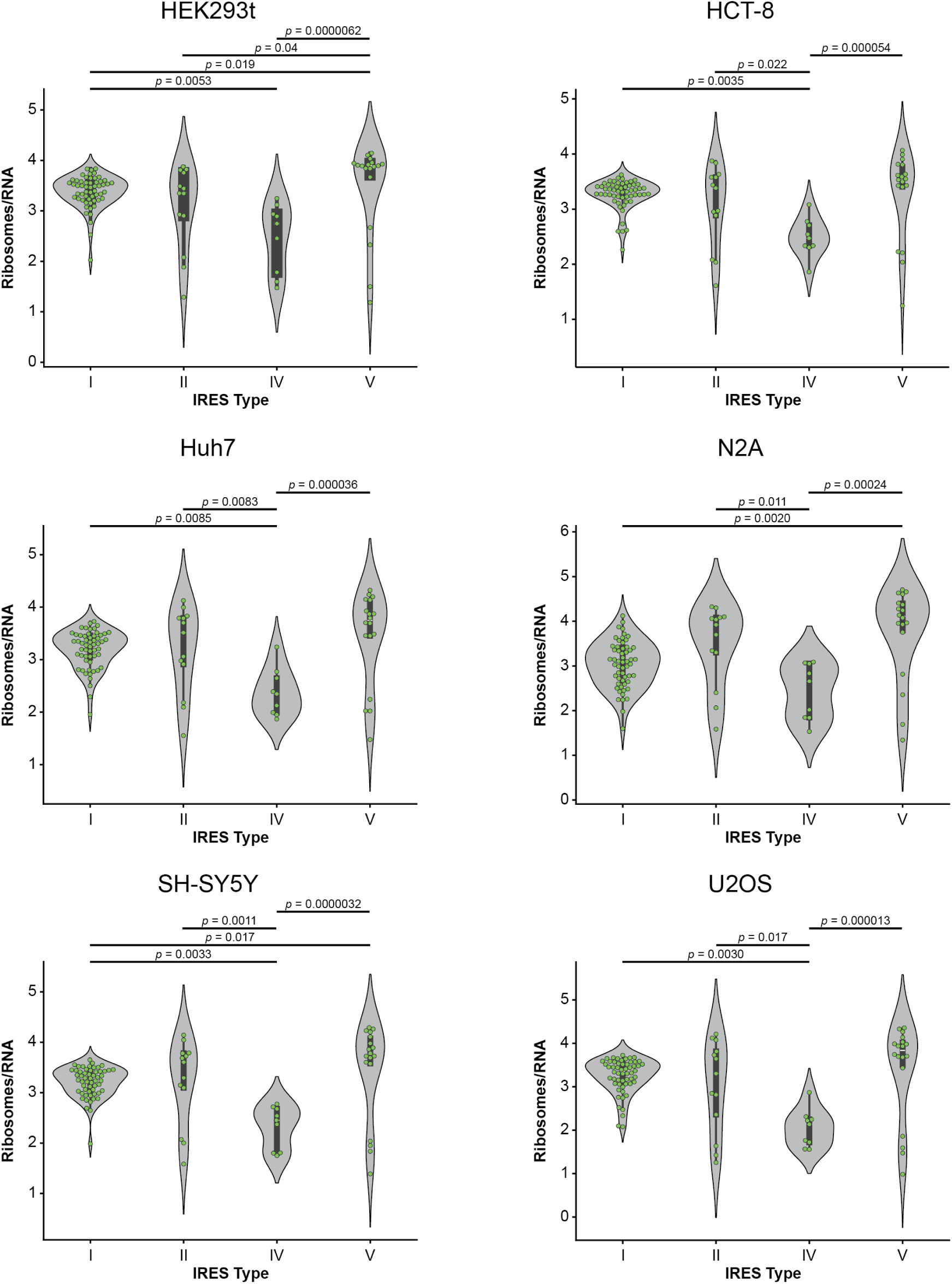
Type V IRESes exhibit high activity across all cell types. Same as Figure 2F, but shown individually for all six cell lines: HEK293T, HCT-8, Huh7, N2A, SH-SY5Y, and U2OS. Violin plots display ribosomes/RNA grouped by IRES structural type (I, II, IV, V). *p*-values from Kruskal-Wallis H test with Dunn posthoc analysis are indicated.

**Figure 6:**
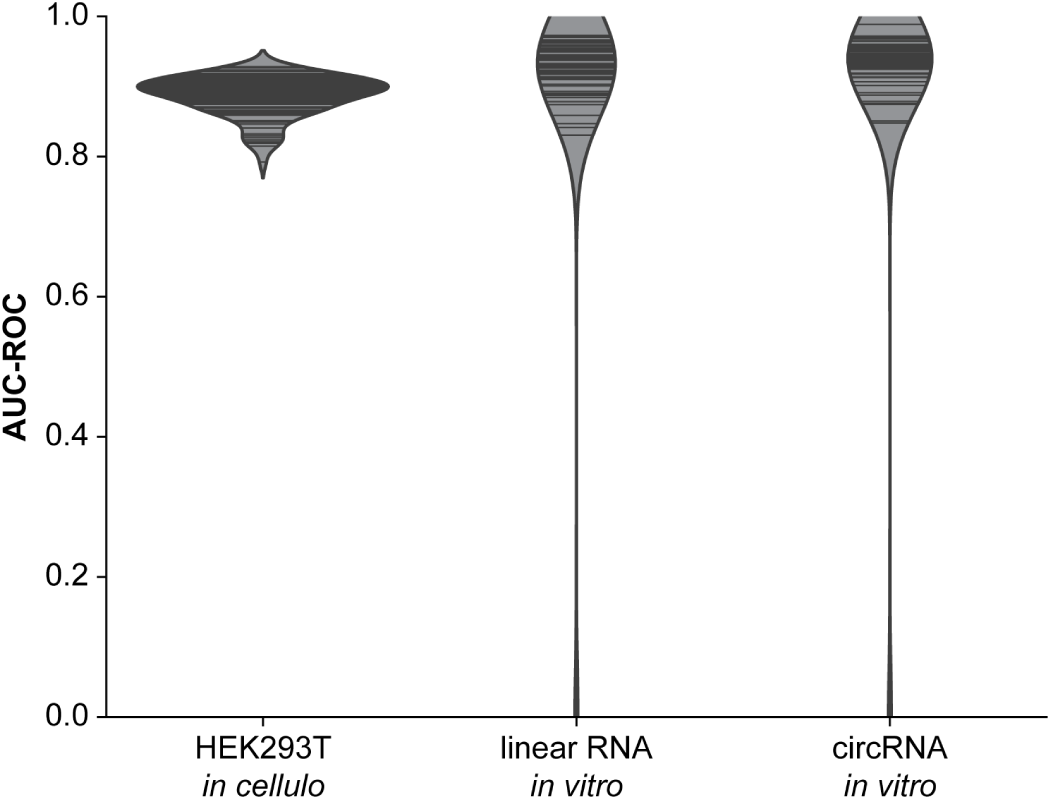
DMS-MaPseq AUC-ROC values in three conditions. AUC-ROC values for three DMS-MaPseq experiments measuring fit of DMS data to the predicting secondary structure model.

**Figure 7:**
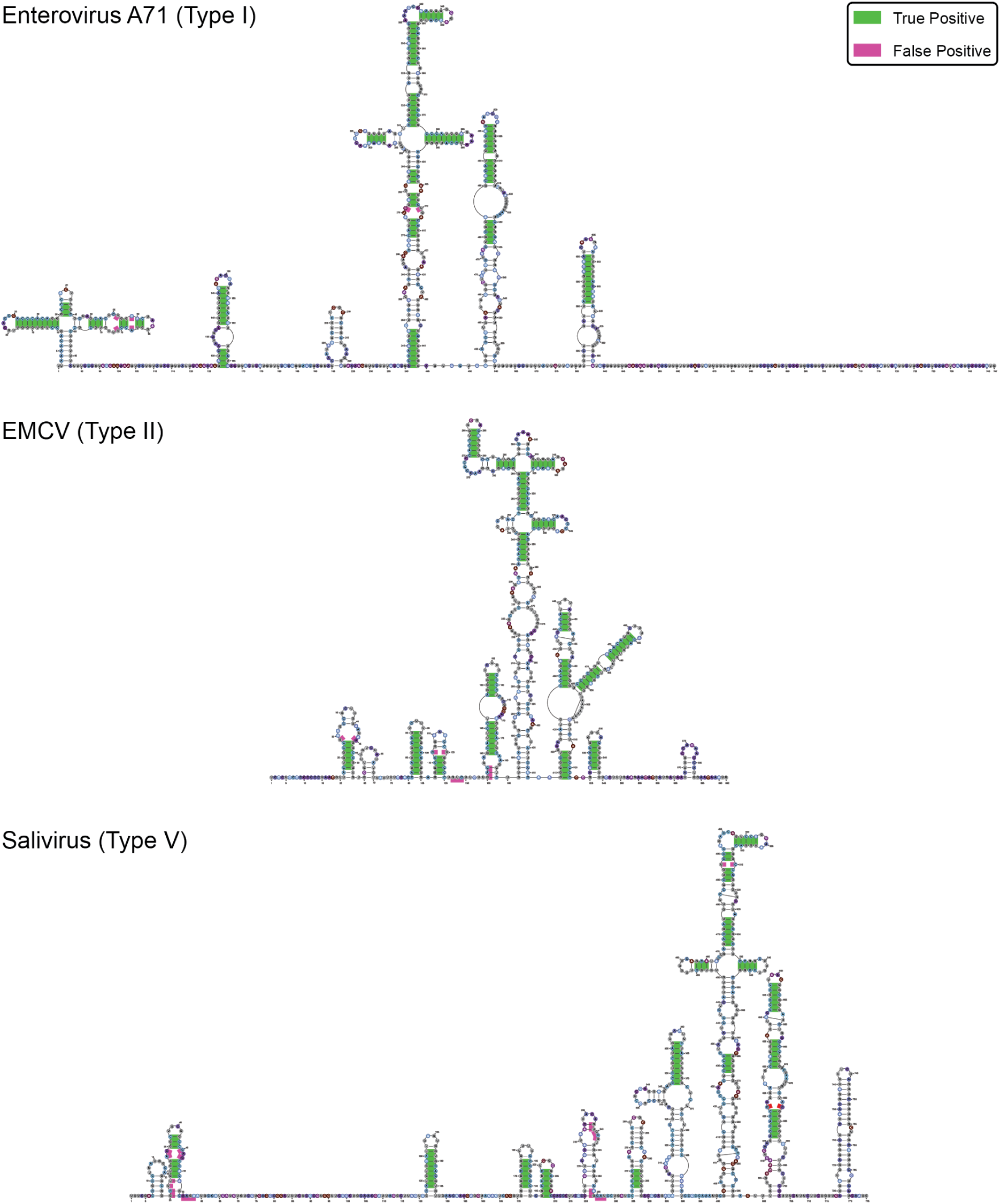
Albatross-predicted base pairs overlaid on consensus literature structures. Representative IRES secondary structures for Enterovirus A71 (Type I), EMCV (Type II), and Salivirus (Type V) are shown. Base pairs predicted by Albatross that agree with the consensus literature structures are colored as true positives (green), while Albatross predictions absent from the consensus literature structures are colored as false positives (pink).

**Figure 8:**
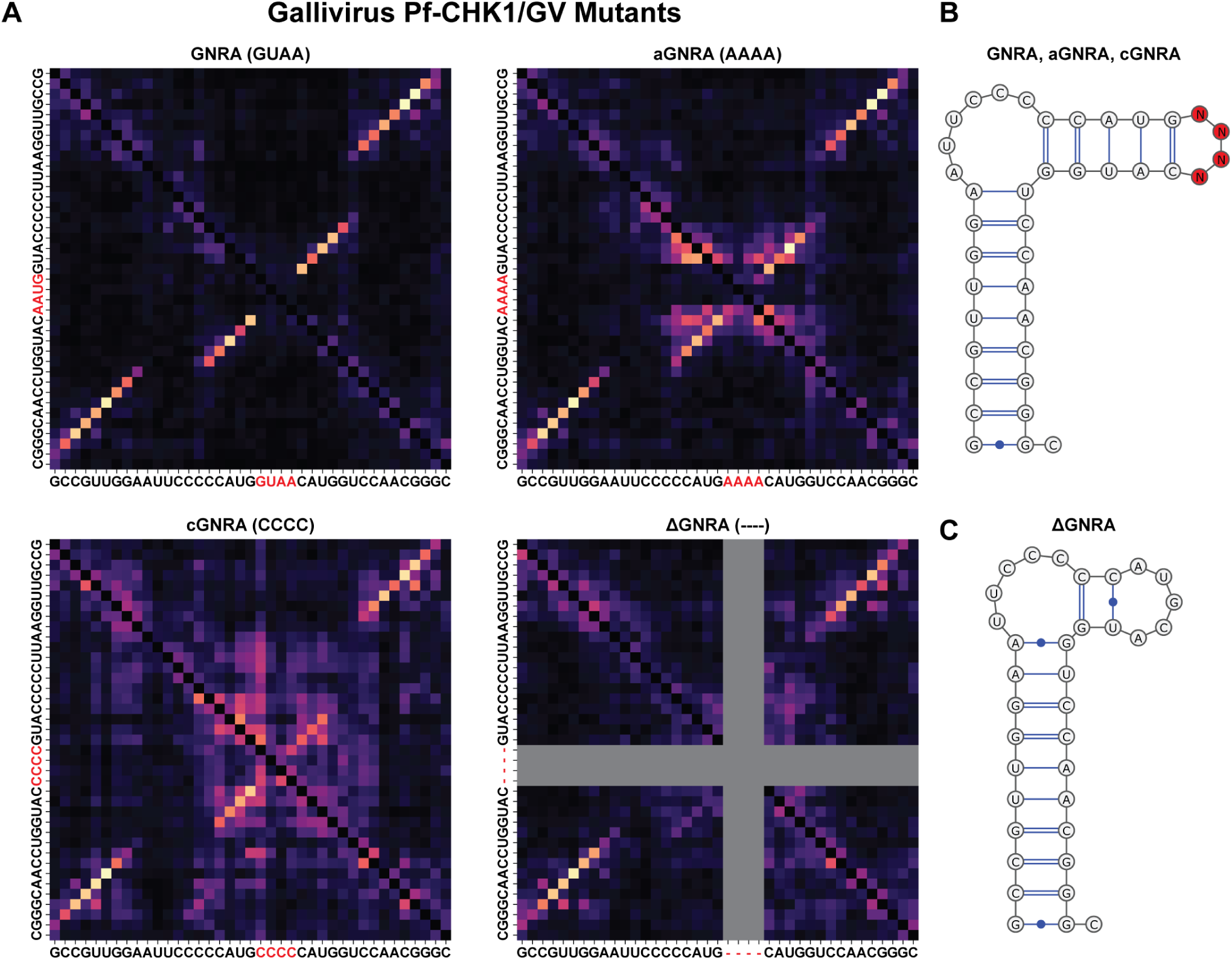
Albatross uses motif recognition to predict structure. (**A**) Dependency maps of the Gallivirus IRES with GNRA mutations: wild-type (GUAA), AAAA, CCCC, and deletion. (**B,C**) Predicted structures for GNRA variants and deletion mutant.

**Figure 9:**
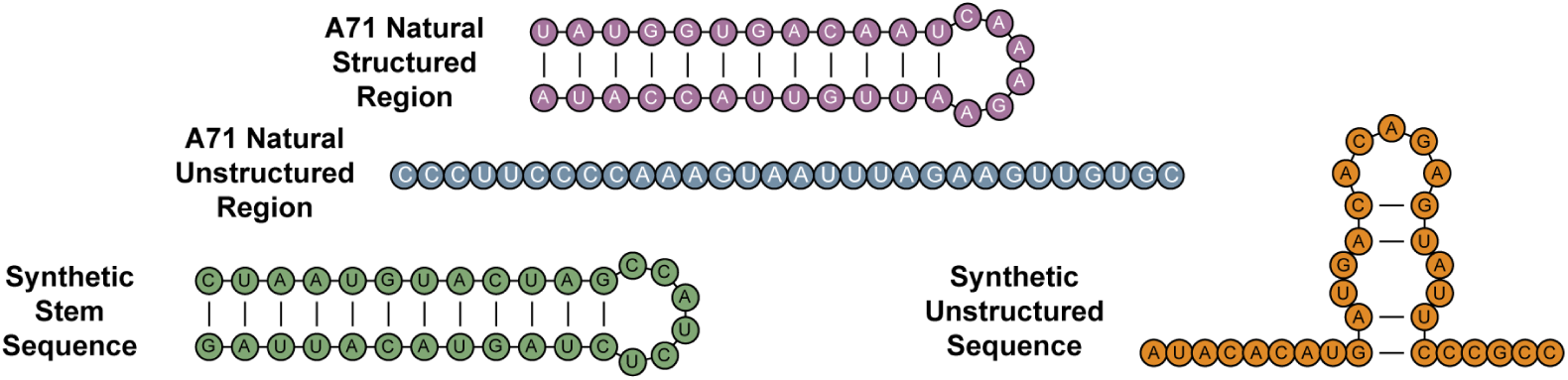
Secondary structures of the four sequence elements for sythnetic sequence prediction experiments. The natural A71 structured region (magenta) and open region (blue), and two synthetic sequences absent from the A71 training set: a thermodynamically stable synthetic stem (green) and a synthetic sequence with low intrinsic folding stability (orange).

**Figure 10:**
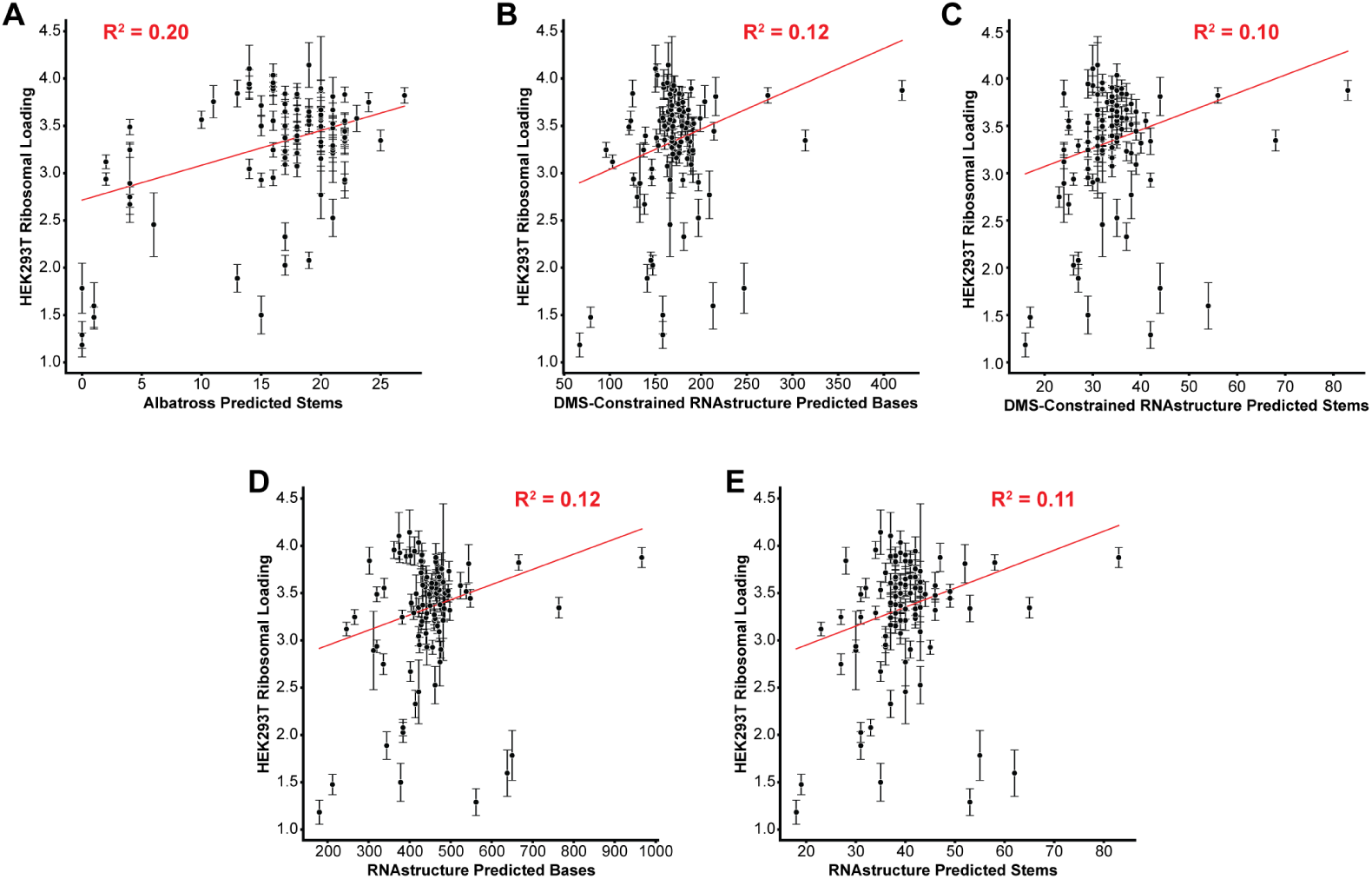
Albatross stem prediction correlates with IRES activity. (**A**) HEK293T activity vs Albatross stem predictions. (**B**) HEK293T activity vs DMS-MaPseq stem predictions. (**C**) HEK293T activity vs RNAstructure stem predictions.

**Figure 11:**
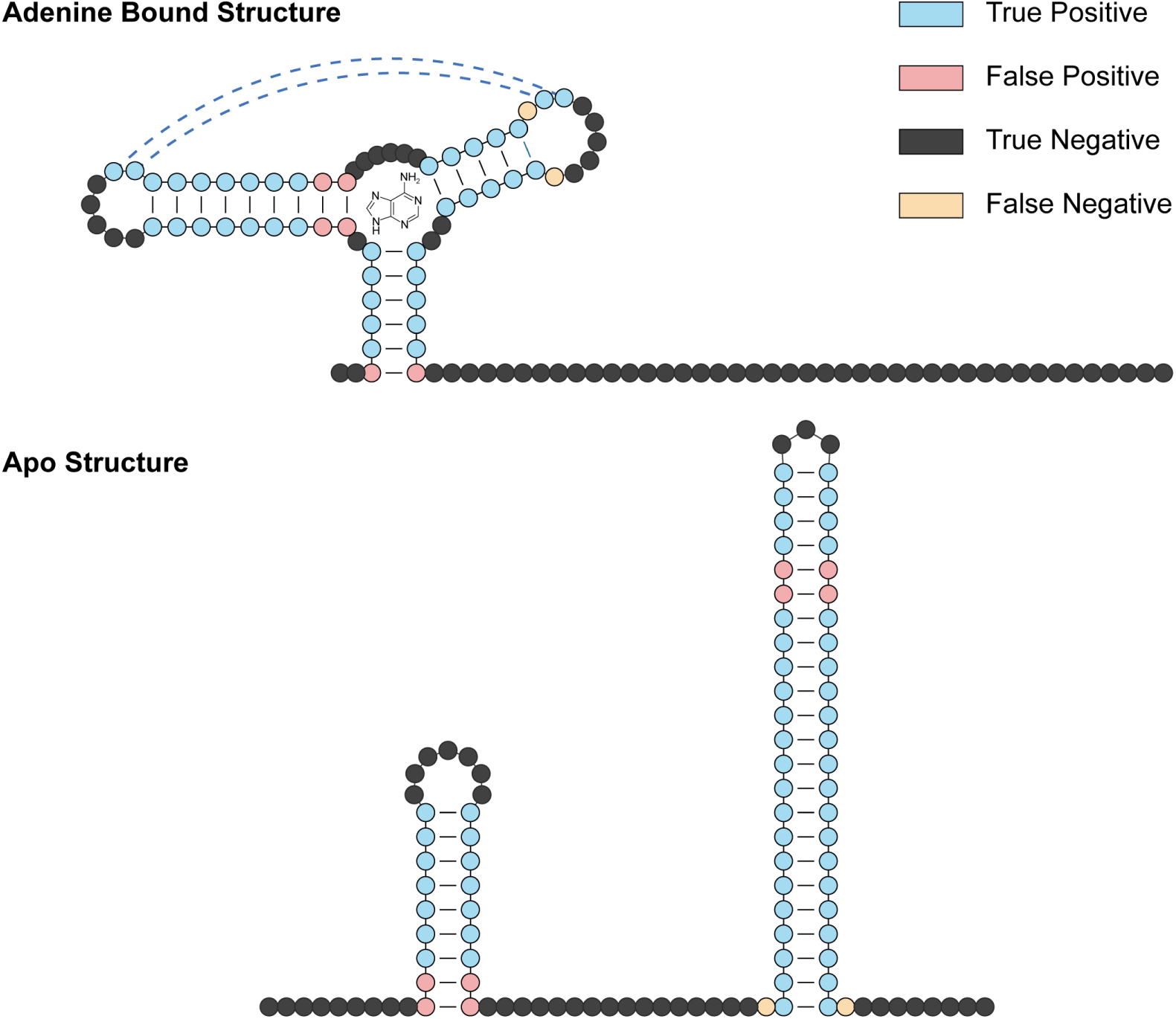
Albatross recovers both conformations of the *B. subtilis* adenine riboswitch. (**Top**) Ligand-bound (holo) structure predicted by Albatross, with the adenine ligand drawn at the three-way junction and the loop-loop interaction indicated by dashed lines. (**Bottom**) Unbound (apo) structure predicted by Albatross. In both panels, nucleotides are colored by agreement with the published structure: true positive (predicted paired and paired in the published structure), false positive (predicted paired, unpaired in the published structure), true negative (predicted unpaired and unpaired), and false negative (predicted unpaired, paired in the published structure). Related to Figure 7G and H.

